# High-throughput microarray analysis provides new insight into optimising Rubisco yield from diverse plants

**DOI:** 10.64898/2026.11.12.766128

**Authors:** Jack C. D. Pattinson, Charlotte Stenkjær Fletcher, Ieva Lelėnaitė, Georg Lietz, William G. T. Willats, Maxim V. Kapralov

## Abstract

Interest in plant-based proteins is increasing due to environmental, ethical, and health considerations. Ribulose-1,5-bisphosphate carboxylase/oxygenase (Rubisco), the most abundant protein on Earth, is present in all green leafy biomass and possesses excellent nutritional and functional properties. Despite its potential, Rubisco remains underutilised because extraction methods are often costly, inefficient, and influenced by species-specific differences. Moreover, the impact on Rubisco yield of cell wall-protein interactions remains poorly understood. We applied Microarray Polymer Profiling (MAPP), a high-throughput platform for simultaneous polysaccharide and protein mapping, to characterise diverse leafy biomass sources. This approach revealed interspecies variation and glycan-protein interactions that influence protein extractability and these insights inform biomass selection and optimisation of extraction protocols to improve Rubisco yield. MAPP was also used to evaluate the effects of carbohydrate-active enzymes on Rubisco recovery through targeted cell wall modification.

**Graphical abstract:** 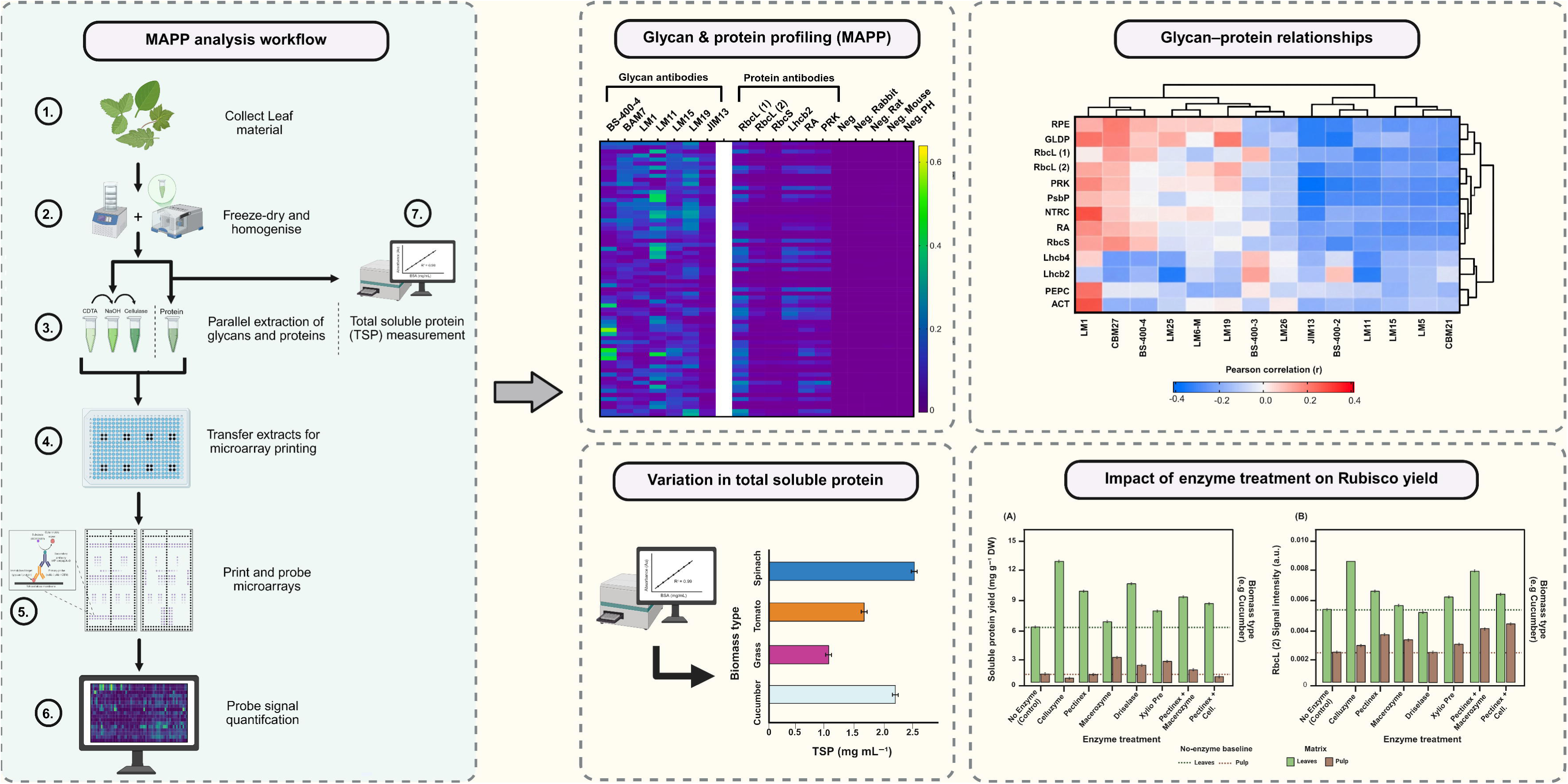

## 1. Introduction

Global agricultural production is facing unprecedented pressure to produce enough nutritious food while simultaneously reducing its environmental impact. With the global population projected to reach approximately 9.7 billion by 2050, rising incomes, urbanisation, and changing dietary preferences are expected to drive greater demand for protein-rich foods (United Nations, Department of Economic and Social Affairs, Population Division, 2022). However, conventional livestock production contributes substantially to greenhouse gas emissions, freshwater use, land degradation, and biodiversity loss (Poore & Nemecek, 2018; Tilman & Clark, 2014; Willett et al., 2019). Consequently, the identification of sustainable protein sources capable of meeting future nutritional requirements without placing additional pressure on natural resources has become a research priority (Aiking & de Boer, 2020; van der Goot et al., 2016).

Plant-derived proteins have emerged as attractive alternatives to animal proteins due to their generally lower environmental footprint and increasing consumer acceptance. Current markets are dominated by soy, pea, and wheat proteins because of their high protein content and established processing technologies (Day, 2013; Kyriakopoulou et al., 2019). However, dependence on a limited number of crop species raises concerns regarding land use, monoculture production, and supply chain resilience. Many conventional plant proteins also require extensive processing to remove antinutritional compounds and undesirable sensory characteristics, increasing production costs and limiting consumer appeal (McClements & Grossmann, 2021). These challenges have stimulated interest in alternative protein sources that can be recovered from underutilised biomass streams while maintaining favourable nutritional and functional properties.

Green leafy biomass represents one of the largest yet least exploited sources of plant protein. Leaves are produced in large quantities as agricultural residues from vegetable, forage, and horticultural crops, while dedicated biomass crops can generate substantial protein yields per hectare (Pérez-Vila et al., 2022; Tamayo Tenorio et al., 2016). Despite this abundance, leafy biomass is commonly used as animal feed, compost, or returned to soil, despite containing large quantities of soluble proteins. Recovering proteins from these biomass streams offers an opportunity to increase the value of existing agricultural systems and supports circular bioeconomy principles aimed at maximising resource efficiency and reducing waste generation (van der Goot et al., 2016).

Among the proteins present in green leaves, ribulose-1,5-bisphosphate carboxylase/oxygenase (Rubisco) has attracted considerable attention because of its abundance, nutritional quality, and functional properties. Rubisco is the key enzyme responsible for carbon fixation during photosynthesis and is widely recognised as the most abundant protein on earth (Andersson & Backlund, 2008; Bar-On & Milo, 2019; Ellis, 1979; Spreitzer & Salvucci, 2002). In many C□ species it accounts for 30-50% of soluble leaf protein and up to 30% of total leaf nitrogen (Bar-On & Milo, 2019; Ellis, 1979; Spreitzer & Salvucci, 2002). This exceptional abundance, combined with the renewability of leaf biomass, makes Rubisco an attractive candidate for sustainable protein production.

Interest in Rubisco has increased as advances in green biorefinery technologies have demonstrated the feasibility of producing food-grade proteins from fresh leaf biomass (Krishnan et al., 2026; Pérez-Vila et al., 2022; Tamayo Tenorio et al., 2016). Unlike seed storage proteins, Rubisco is recovered from vegetative tissues that are continuously produced throughout plant growth and are often underutilised. Exploiting Rubisco therefore offers the potential to increase protein production without requiring additional agricultural land or major changes to existing cropping systems. Moreover, Rubisco possesses several characteristics that make it attractive as a food ingredient, and it is considered one of the highest-quality plant proteins available. Unlike many seed storage proteins, which evolved primarily as nutrient reserves, Rubisco functions as a metabolic enzyme and exhibits a balanced amino acid composition that closely aligns with human nutritional requirements (Shewry, 2007). It contains high concentrations of lysine, threonine, valine, leucine, and isoleucine, amino acids that are often limiting in cereal proteins (Rhodes et al., 1980; Tamayo Tenorio et al., 2016). Furthermore, purified Rubisco preparations are pale in colour, possess a relatively neutral flavour and contain relatively low levels of antinutritional compounds such as trypsin inhibitors and lectins (Food and Agriculture Organization of the United Nations, 2013; Martin et al., 2019; McClements & Grossmann, 2021). Previous studies have demonstrated excellent solubility across a wide pH range, together with strong emulsifying, foaming, and gel-forming capacities (Krishnan et al., 2026; Ma et al., 2022; Pérez-Vila et al., 2022; Tamayo Tenorio et al., 2016). These characteristics make Rubisco a promising ingredient for a variety of food applications, including beverages, dairy alternatives, aerated products, and meat analogues. Compared with conventional plant proteins, Rubisco offers a unique combination of nutritional quality and functional performance, suggesting it could complement existing protein ingredients in future food systems. Despite these advantages, commercial utilisation of Rubisco remains limited by challenges associated with efficient extraction and purification.

Rubisco is located within the chloroplast stroma and is enclosed by multiple structural barriers, including chloroplast membranes, the plasma membrane, and the surrounding plant cell wall (Pérez-Vila et al., 2022; Tamayo Tenorio et al., 2016). Conventional extraction methods generally involve mechanical disruption of fresh biomass followed by pressing and separation of protein-rich green juice from fibrous residues. However, these approaches often recover only a fraction of the soluble protein present within leaf tissues, with substantial quantities remaining associated with the fibre fraction (Pérez-Vila et al., 2022; Tamayo Tenorio et al., 2016). In addition, extraction efficiency varies considerably among plant species, suggesting that intrinsic biological characteristics strongly influence protein accessibility and release.

The plant cell wall is increasingly recognised as a major barrier limiting recovery of intracellular proteins (Kleuter et al., 2024; Sharma et al., 2025; Wang et al., 2025). Cell walls are complex composite structures composed primarily of cellulose, hemicelluloses, pectin, structural proteins, and phenolic polymers (Carpita & Gibeaut, 1993; Cosgrove, 2005; Cosgrove, 2018; Harholt et al., 2010; Ridley et al., 2001; Scheller & Ulvskov, 2010; Showalter, 1993; Somerville et al., 2004). Their composition and organisation vary according to species, tissue type, and developmental stage, influencing wall rigidity, porosity, and susceptibility to disruption (Carpita, 2011; Fangel et al., 2012). Such variation is likely to affect the release of Rubisco during processing and hence the effectiveness of extraction technologies. However, while considerable effort has been directed towards optimising extraction parameters such as pH, temperature, and mechanical pressing, relatively little attention has been given to understanding how differences in cell wall composition influence protein recovery (Pérez-Vila et al., 2022; Tamayo Tenorio et al., 2016).

One promising approach to overcoming these structural barriers is enzyme-assisted extraction. Cellulases, hemicellulases, and pectinases selectively degrade major components of the cell wall, increase tissue permeability and enhance access to intracellular proteins (Harholt et al., 2010; Scheller & Ulvskov, 2010). Previous studies have shown that enzyme-assisted processing can improve protein recovery and reduce the energy required for biomass disruption (Krishnan et al., 2026; Ma et al., 2022; Pérez-Vila et al., 2022; Tamayo Tenorio et al., 2016). However, responses vary substantially among species, suggesting that differences in cell wall architecture influence the effectiveness of enzyme treatments and the specific relationships between cell wall polysaccharide composition and Rubisco extractability remain poorly understood.

In this study, we used an analytical platform (Microarray Polymer Profiling or ‘MAPP’) that combines the specificity of molecular probes (monoclonal antibodies, mAbs, and carbohydrate binding modules, CBMs) with the high-throughput capacity of microarrays (Knox, 2008; Møller et al., 2007; Pattathil et al., 2010; Pattathil et al., 2015; Pedersen et al., 2012; Verhertbruggen et al., 2009). We assessed Rubisco abundance in 128 phylogenetically diverse plant species and investigated plant cell wall polysaccharides that may influence biomass processing and Rubisco extraction. By linking variations in cell wall architecture with protein recovery, we aimed to identify the key cell wall structural determinants limiting Rubisco extraction and to establish new principles for feedstock selection and process optimisation. These insights address a critical barrier to the commercial utilisation of leaf proteins and provide a framework for developing more efficient, species-specific extraction technologies.

## 2. Materials and methods

All chemicals and reagents were of analytical grade, unless otherwise stated.

### 2.1 Collection of plant material

Leaves from a representative sample set of 128 plants were used to understand the impact of cell wall polysaccharides on soluble leaf protein Rubisco extractability. The plants chosen were a representation of common crops, house and commonly encountered garden plants, where plant material was obtained from living plants. The species in the collection are available at Newcastle University. Details of the plant material used are included in **Table S1.**

### 2.2 Sample preparation

For each species, leaves were collected in abundance from multiple plants where possible. The leaves were combined into a Falcon tube for each species to normalise sampling per species and immediately flash-frozen in liquid nitrogen. The samples were then stored in the -80 °C before being freeze-dried (LaboGene). The freeze-dried leaves were homogenised by shaking using one 12 mm steel bead per sample. Once homogenised, 3 samples of approximately 30 mg, weighed with 1 decimal, was taken from each Falcon tube to yield 3 replicate samples per species. This procedure was repeated to yield a total of 6 replicate samples per species to account for the two-part extraction series.

### 2.3 Protein and polysaccharide extractions

A modified two-part extraction series was used to extract constituent glycans and proteins. The extractant volumes were adjusted to the exact weight of each replicate sample (30 µL mg^-1^). The extraction series is based on the procedure used by Bakshani et al. (2023) and Møller et al. (2007) with the addition of a cellulase extraction and a separate protein extraction.

### 2.4 Protein extraction

Plant leaf material has a large amount of protein present, hence the first extraction targeted these. 3 replicate samples per species were reconstituted in a 50 mM HEPES, pH 7.7, 150 mM NaCl, 1 mM EDTA, 2 mM MgCl_2_, 10 % (v/v) Glycerol, 0.1 % Triton X-100, 1 % (w/v) PVPP and 1 µL mL^-1^ of protease inhibitor cocktail (added just before use). A steel bead was added to each sample tube and then shaken at 27 s^-1^ for 2 minutes in a Tissuelyser II (Qiagen, UK). The speed was then reduced to 10 s^-1^ for 2 hours. After extraction, the samples were centrifuged at 13000 RPM for 10 minutes (min). The supernatant was carefully removed from each of the samples and transferred into an Eppendorf. Extracts were stored at 4 °C.

### 2.5 Polysaccharide extraction

A new set of 3 replicate samples per species were subjected to a 3-step sequential extraction. Firstly, the solvent 50 mM CDTA (pH 7.5) was used, a steel bead was added, and the extraction process was carried out as described above. The second extraction step used a 4 M NaOH + 0.1 % (w/v) NaBH_4_ buffer and the extraction step was similarly carried out as described above. The third and final extraction of the precipitate pellet from second extraction step used 20 mM Tris, pH 8.8, with the addition of 2 µL mL^-1^ of *Bacillus sp.* Cellulase 5A (NZYTech, Portugal). This extraction buffer requires a 16-hour incubation at 45 °C, on a dry heat block followed by centrifugation at 13000 RPM for 10 min, and the supernatant being carefully removed. All extracts were stored at 4 °C.

### 2.6 Microarray printing

A serial dilution of each sample was made in a 384-well microtiter plate. Each extract was diluted 4-fold in ArrayJet buffer (47 % glycerol, 52.9 % deionised water, 0.06 % Triton X-100 and 0.04 % ProClin™ 200) to a final volume of 40 μL in each well. Plates with diluted extracts were centrifuged at 3700 RPM for 10 min before they were printed on a nitrocellulose membrane using an Arrayjet Marathon Argus Microarray Printer (Arrayjet, Roslin, UK) as described previously (Bakshani et al., 2023; Fangel et al., 2024). For each sample, the dilution series was printed in duplicate on each microarray.

### 2.7 Microarray probing

The printed arrays were probed and developed following procedures described by Bakshani et al. (2023) and Fangel et al. (2024). A total of 53 arrays were used for this analysis, one for each antibody and 5 for negative controls of the secondary antibodies. They were cut and placed into 4-well plate and a 5 % (w/v) semi-skimmed milk powder solution (MPBS) was made in phosphate-buffered saline (PBS; 140 mM NaCL, 2.7 mM KCl, 10 mM Na_2_HPO_4_, 1.7 mM KH_2_PO_4_, pH 7.5). The arrays were left agitating in MPBS to block any non-specific binding to the nitrocellulose for 1 hour. The arrays were then placed face down onto an MPBS solution containing one of the antibodies; one array is used per primary antibody. The control arrays were left in the pure MPBS solution because they are needed for negative controls of the secondary antibodies. The primary antibody solutions were made by adding antibodies to the MPBS solution and aspirating thoroughly. The optimal primary antibody concentrations were given by the manufacturers (**Table S2**). The microarrays were left in the antibody and MPBS solution for 2 hours and then washed in PBS 4 times and left agitating for 5 minutes after every 2 washes. The microarrays were subsequently probed with the secondary antibody, either anti-rat, anti-mouse, anti-rabbit or anti-polyhistidine, depending on the origin of the primary antibody used on the specific array (**Table S2**). The secondary antibodies were conjugated to alkaline-phosphatase (Jackson ImmunoResearch and Sigma, Merck Life Science) and diluted 1:1000 in MPBS. The arrays were left agitating in the secondary antibody and MPBS solution for 2 hours before being washed in PBS as described above. Finally, the arrays were submerged in a developing solution containing 5-bromo,4-chloro,3-indolylphosphate (BCIP) and 4-Nitro blue tetrazolium chloride (NBT). BCIP/NBT was added to an alkaline phosphatase buffer (100 mM Tris, 5 mM MgCl_2_•6H_2_O, pH 9.5). The developing arrays were rinsed in deionised water before they were mounted on a thin sheet of commercially available graph paper and prepared for scanning.

### 2.8 Microarray quantification

The developed arrays were scanned with a Canon 9000F Mark II desktop scanner (Canon, Newcastle, England) using IJ Scan Utility at a resolution of 2400 dpi. The arrays were scanned individually using grayscale colour settings, and the images were saved as JPG files. The antibody binding signals are visualised on the arrays as lighter or darker spots, thus meaning darker spots present higher concentrations of a specific epitope. If a specific epitope is not present in a sample, there will be no binding of the antibody, hence no spot is shown on the grayscale array image.

The array images were uploaded and quantified one at a time using ArrayPro Analyzer (v.6.3.1, Media Cybernetics, Rockville, USA). Spots were quantified by overlaying a unique .GAL file, and spots are defined as the mean pixel value within each spot minus the median pixel value of the surrounding local background. The data were then exported and saved as a .txt file and analysed using Microsoft Office Excel (Microsoft, Redmond, WA). Averages were calculated for each set of replicates across each of the dilutions and the highest value of the data set was assigned the value of 100. The rest of the data set was adjusted accordingly as described by Bakshani et al. (2023) and Fangel et al. (2024).

### 2.9 Total soluble protein (TSP) quantification

Total soluble protein (TSP) concentrations (mg mL□¹) were determined using the Pierce Bradford Protein Assay Kit (Thermo Fisher Scientific; catalogue no. 23200), following the manufacturer’s microplate protocol. The assay is based on the Bradford method (Bradford, 1976), and absorbance was measured using a Varioskan LUX multimode microplate reader (Thermo Fisher Scientific, Waltham, MA, USA).

### 2.10 Enzyme Treatment

Enzyme treatments were conducted using an adaptation of Vidal-Melgosa et al. (2015). Leaf material from cucumber (*Cucumis sativus*), tomato (*Solanum lycopersicum*) and perennial ryegrass (*Lolium perenne*) was obtained from the same biomass batches used for the screening panel. Pulp was prepared by thawing 200 g of frozen leaves at 4 °C and processing them using a twin-screw press (Angel Juicer 5500, Angel Co., Ltd., Busan, Republic of Korea); the resulting solid fraction was retained. Freeze-dried and frozen-thawed leaf and pulp samples were prepared in triplicate. Freeze-dried material was weighed directly to approximately 30 mg dry weight (DW), whereas frozen-thawed material was adjusted using its measured dry matter content to provide an equivalent dry mass (Table S3). Seven enzyme treatments were evaluated: Celluzyme 0.7T, Pectinex Ultra SP-L, Macerozyme R-10, Driselase, Xylio Pre, Pectinex Ultra SP-L combined with Macerozyme R-10, and Pectinex Ultra SP-L combined with Celluzyme 0.7T. Samples were suspended in 50 mM sodium acetate buffer (pH 5.5) containing the relevant enzyme preparation at 0.1–0.5 % (w/v), or buffer alone as the no enzyme control. Treatment specific enzyme concentrations are provided in Table S3. Samples were incubated at 30 °C for 1 hour with gentle agitation. A steel bead was then added, and the samples were disrupted at 10 s□¹ for 1 hour using a TissueLyser II (Qiagen, UK), followed by centrifugation at 13000 RPM for 10 min. Supernatants were stored at 4 °C until MAPP (**section 2.6 - 2.8**) and TSP (**section 2.9**) analyses, was carried out. Please note, only RbcL (2) was used for probing.

### 2.11 Data preparation and visualisation

Unless otherwise stated, measurements were performed using three replicates and are presented as mean ± SD. Replicates were averaged for the biomass-screening analyses, whereas enzyme-treatment analyses used replicate-level data. Enzyme treatment TSP yield was expressed as mg g□¹ DW and calculated by multiplying TSP concentration by extraction volume and dividing by sample dry mass. The weighed mass was used for freeze-dried material, while frozen-thawed dry mass was calculated from the wet mass and measured dry matter fraction (**Table S3**). Protein and glycan antibody heatmaps were generated in GraphPad Prism (v11.0.2; GraphPad Software, San Diego, CA, USA); all other figures were generated in R using RStudio.

### 2.12 Statistical analysis

Statistical analyses were performed using R v4.5.2 (R Core Team, 2025) within RStudio v2026.07.1+147. Associations between TSP and Rubisco antibody signals were assessed using two-sided Pearson correlations and ordinary least-squares regression with 95% confidence intervals. Protein-glycan associations were evaluated using pairwise-complete Pearson correlations between signals from 13 protein and 15 glycan antibodies across 128 biomass samples (195 comparisons). Associated p values were adjusted across the 195 comparisons using the Benjamini-Hochberg false discovery rate. Correlation profiles were hierarchically clustered using Euclidean distance and complete linkage. TSP was included separately in a sensitivity analysis (**Fig. S1**).

Principal component analyses used complete, mean-centred and unit-variance-scaled data. PC1 scores and loadings were jointly reversed where necessary so that the summed loadings of the three Rubisco antibodies were positive. Ellipses drawn at the 68% level summarised within-group dispersion. Group effects were assessed on the complete standardised matrices using permutational multivariate analysis of variance (PERMANOVA) with Euclidean distances and 9,999 permutations. Differences in dispersion were assessed using permutational analysis of multivariate dispersion (PERMDISP). The vine/climber category (n = 2) was excluded from the primary growth-form tests because the group size was insufficient for stable inference; including all groups in a sensitivity analysis did not alter the overall interpretation. Associations between the six targeted glycan antibodies and the multivariate response across the three Rubisco antibodies were assessed by redundancy analysis (RDA) using permutation testing and adjusted R². Conditional associations with Rubisco-only PC1 and TSP were examined using standardised multiple linear regression with HC3 robust standard errors and 95 % confidence intervals. Pearson and Spearman correlations were used as sensitivity analyses, with Benjamini-Hochberg adjustment applied within each analysis family.

Enzyme-treatment effects on TSP yield and RbcL (2) signal in frozen-thawed material were analysed separately within each species × matrix combination by one-way analysis of variance (ANOVA), followed by Dunnett-adjusted comparisons of each treatment with the matched no enzyme control. Model assumptions were assessed using residual diagnostic plots, Shapiro-Wilk tests for residual normality and Levene tests for homogeneity of variance. Unless otherwise stated, statistical significance was accepted at *p* < 0.05.

## 3. Results and discussion

### 3.1 Development of a high-throughput analytical platform

Using MAPP technology as a basis, we developed a specialised platform for the high-throughput parallel analysis of proteins involved in photosynthesis (including Rubisco) and polysaccharides from 128 samples of plant biomass. The workflow used is shown **Fig. 1** and involved extraction of polysaccharides and proteins which were printed as microarrays and subsequentially interrogated using panels of antibodies (**Table S2**). Microarray data is presented as heatmaps (full heatmap can be seen in **Data S1**). TSP was also determined using Bradford assays from the same sample set. It is important to recognise that MAPP data does not provide information about absolute amounts of glycans or proteins but rather reports differences in relative abundance across samples.

**Figure 1.**
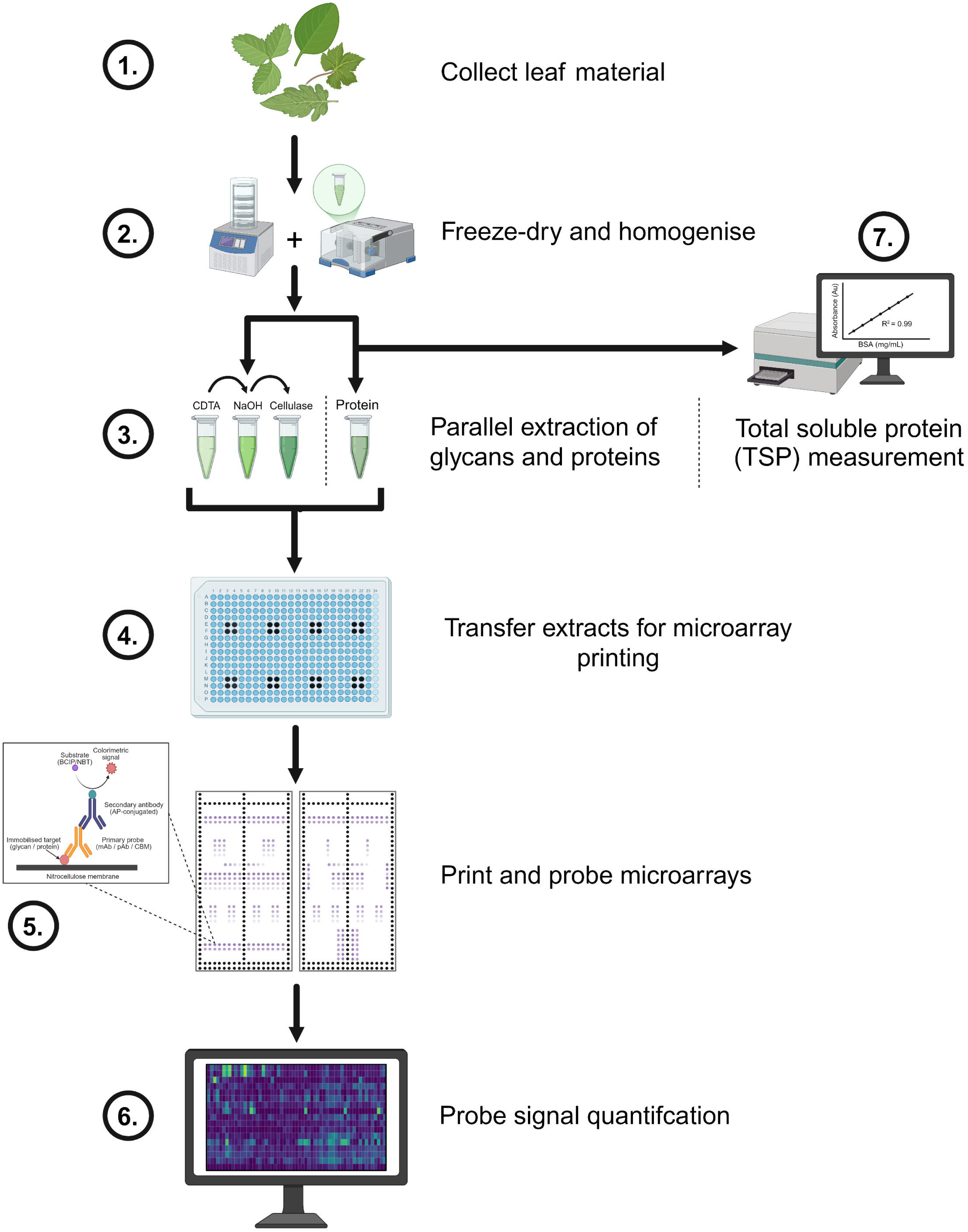
Overview of the experimental workflow used for parallel glycan and protein profiling of leaf biomass. Leaf material was (1) collected and (2) freeze-dried and homogenised. (3) Glycans and proteins were extracted in parallel using the indicated extraction procedures. (4) The extracts were transferred to microplates for microarray printing, and (5) the printed arrays were probed with selected glycan- and protein-binding reagents. (6) Probe-binding signals were quantified and visualised for subsequent analysis. In parallel, (7) protein extracts were analysed for total soluble protein (TSP). This workflow enabled integrated comparison of cell-wall glycan epitope profiles, protein-probe signals and TSP concentrations across the biomass panel.

### 3.2 MAPP analysis of Rubisco and other photosynthesis related proteins

MAPP was used to assess the relative abundance of Rubisco and associated photosynthetic proteins across the plant sample set. Microarrays of extracted proteins were probed with monoclonal antibodies specific for Rubisco subunits and a variety of other protein with data presented as a heatmap in **Fig. 2**.

**Figure 2.**
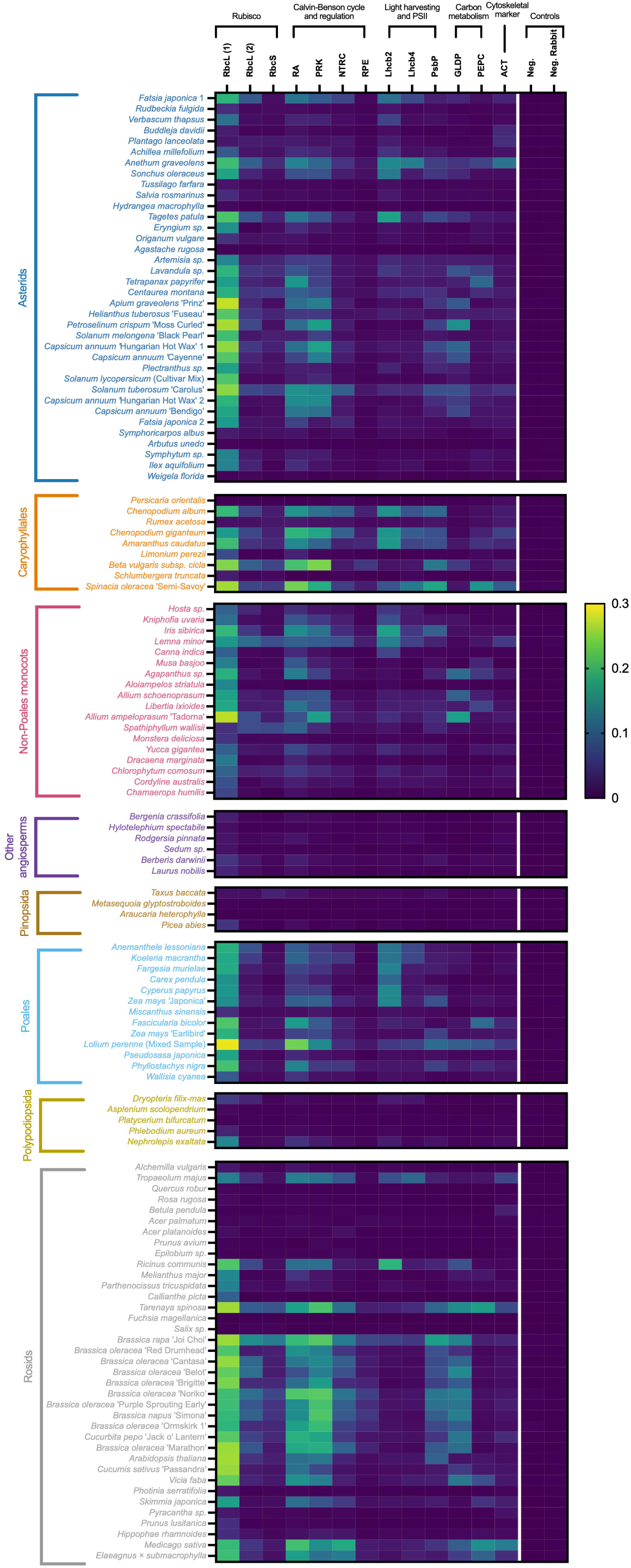
Protein antibody binding profiles across the biomass panel. Rows represent biomass samples and columns represent functionally grouped protein antibodies; relative signal intensity (0–0.3) increases from blue through green to yellow. The antibodies targeted the Rubisco large subunit [RbcL (1), Abcam antibody; RbcL (2), Agrisera antibody], Rubisco small subunit (RbcS), Rubisco activase (RA), phosphoribulokinase (PRK), NADPH-dependent thioredoxin reductase C (NTRC), ribulose-phosphate 3-epimerase (RPE), LHCII type II chlorophyll a/b-binding protein (Lhcb2), CP29 chlorophyll a/b-binding protein of photosystem II (Lhcb4), oxygen-evolving enhancer protein 2 of photosystem II (PsbP), glycine decarboxylase P-protein (GLDP), phosphoenolpyruvate carboxylase (PEPC) and actin (ACT). Controls comprised a general negative control (Neg.) and a rabbit secondary-antibody control (Neg. Rabbit). Antibody specificities, dilutions, suppliers and references are provided in Supplementary table 2.

The relative abundance of extracted proteins varied widely across the biomass sample set, as reported by different binding signals. Regarding the three Rubisco specific antibodies (anti-RBcL (1), anti-RbcL (2) and RbcS), *Brassica rapa* ‘Joi Choi’ produced the highest RbcL (2) and RbcS signals, while *Lolium perenne* produced the highest RbcL (1) signal. *Tarenaya spinosa*, *Beta vulgaris* subsp. *cicla* and *Spinacia oleracea* ‘Semi-Savoy’ also showed consistently high Rubisco related profiles. Conversely, *Fuchsia magellanica* and *Araucaria heterophylla* showed lower signals for all three probes. RbcL (1/2) and RbcS were the most strongly associated antibodies in terms of binding profiles, consistent with their stoichiometric assembly in the Rubisco holoenzyme and coordinated expression across the chloroplast and nucleus (Suzuki & Makino, 2012; Vitlin Gruber & Feiz, 2018). However, there were also notable variations in the binding of individual Rubisco antibodies within some biomass samples. For example, *Anemanthele lessoniana* produced a high RbcL (2) signal but a substantially lower RbcS signal, whereas *Pelvetia canaliculata* had low RbcL (1) and higher RbcL (2)/RbcS signals. This supports the notion that RbcL and RbcS protein concentration is not necessarily present as a 1:1 ratio, and as such must be measured separately to understand the abundance of each (Recuenco-Muñoz et al., 2015; Wostrikoff & Stern, 2007). These findings also highlight the importance of using a panel of three antibodies with specificity for different subunits to obtain a more complete picture of overall Rubisco abundance. Signals from Calvin–Benson cycle/regulatory proteins, photosystem II targets and associated carbon-metabolism enzymes were also heterogeneous and did not completely parallel the Rubisco profiles, indicating that the array resolved complementary components of the photosynthetic-protein phenotype.

Antibody binding profiles were not generally correlated to taxonomic groups, with Joi Choi, Tarenaya, sugar beet, spinach and Lolium yielding high signals despite their phylogenetic divergence. This is consistent with evidence that nitrogen allocation to Rubisco varies with leaf traits and environment rather than following a single plant-functional-type rule (Luo et al., 2021). Developmental state also affects both the soluble-protein pool and extractability, and a recent tomato leaf proteomics study linked declining extractability during maturation to a shift from anabolic to catabolic/stress functions and to incomplete recovery of soluble proteins (Kleuter et al., 2025). These data also highlight the importance of empirical phenotyping of candidate feedstocks instead of assuming that crop identity or taxonomy predicts Rubisco potential.

Plants are classified according to their photosynthetic carbon fixation pathway into C_3_, C_4_ and crassulacean acid metabolism (CAM) plants (**Data S2)**. C_3_ plants, which include most temperate crops and dicotyledonous species such as tomato (*Solanum lycopersicum*) and cucumber (*Cucumis sativus*), fix atmospheric CO□ directly through the Calvin–Benson cycle via Rubisco, producing the three-carbon compound 3-phosphoglycerate as the first stable product (Taiz et al., 2018). In contrast, C_4_ plants, including grasses such as maize and many forage species, initially fix CO□ into four-carbon compounds using phosphoenolpyruvate carboxylase (PEPC) before concentrating CO□ around Rubisco within specialised bundle sheath cells. This spatial separation reduces photorespiration and improves photosynthetic efficiency under conditions of high temperature, intense light and limited water availability (Hatch, 1987; Sage & Zhu, 2011). CAM plants, which include many succulents and cacti, separate carbon fixation temporally by opening their stomata at night to minimise water loss while storing CO□ as organic acids that are subsequently decarboxylated during daylight for photosynthesis. This adaptation provides exceptional water-use efficiency in arid environments (Winter & Smith, 2021). The differing photosynthetic pathways are associated with distinct leaf anatomies, physiological characteristics and carbon allocation patterns, all of which may influence biomass composition and the efficiency of protein extraction from plant tissues. In this study, as previously described, C_3_ plants demonstrate higher Rubisco content whilst C_4_ demonstrate lower Rubisco content (Galmés et al., 2014; Sugawara et al., 2026).

MAPP highlighted key differences in plant protein abundance across the sample set, demonstrating the effectiveness of this technique as a way to screen a large sample set in a high-throughput manner. Next, we assessed the extent to which MAPP data agreed with those generated using the established Bradford assays to measure TSP.

### 3.3 TSP – Variation in extractable soluble protein across the biomass panel

The Bradford assay has been used for decades to quantify TSP due to its simplicity, rapidity, cost-effectiveness, and relative sensitivity. In the context of measuring Rubisco across the sample set, Bradford measurements are informative because Rubisco often constitutes a substantial proportion of soluble leaf protein.

Extractable soluble protein was quantified across the biomass panel using a standardised protein extraction workflow (**section 2.4**). TSP concentrations varied markedly among the 128 samples (**Fig. 3**), ranging from 0 to 2.906 mg mL□¹, with a median of 0.423 mg mL□¹ and an interquartile range of 0.083–1.060 mg mL □¹. *Spinacia oleracea* ‘Semi-Savoy’ produced the highest concentration (2.906 ± 0.032 mg mL□¹), followed by *Brassica rapa* ‘Joi Choi’ (2.662 ± 0.130 mg mL□¹), *Tarenaya spinosa* (2.310 ± 0.253 mg mL□¹), *Lemna minor* (2.211 ± 0.161 mg mL□¹) and *Chenopodium giganteum* (2.147 ± 0.032 mg mL□¹). High-TSP samples occurred across several taxonomic groups, whereas near-baseline recovery occurred among several fern, conifer and woody ornamental samples. Caryophyllales and rosids contained several highly ranked samples, however, unequal taxonomic representation and substantial within-group variation preclude conclusions relating TSP to plant lineage. This broad variation is consistent with reported crop- and tissue-dependent differences in TSP. Song et al. (2015) identified comparatively high TSP in broccoli and Chinese cabbage, broadly agreeing with the presence of six *Brassica* accessions among the 20 highest-ranked samples in the present screen. However, their use of different plant organs, extraction conditions and a fresh-biomass reporting basis preclude direct value comparison. Among the horticultural crops central to the wider study, spinach produced 2.906 mg mL□¹ TSP, compared with 0.974 mg mL□¹ for cucumber and 0.560 mg mL□¹ for tomato. Spinach therefore showed approximately 3-fold more TSP than cucumber and 5.2-fold more than tomato, consistent with previous studies highlighting spinach as a promising source material for Rubisco extraction and leaf protein concentrate production (Valente et al., 2022). The high ranking of duckweed (*L. minor)* likewise agrees with its recognised potential as a source of Rubisco-rich protein concentrates (Nieuwland et al., 2021). Comparable food- or feed-directed evidence for protein recovery from cucumber leaves remains limited. The intermediate cucumber TSP concentration should therefore be interpreted as a screening-specific indication of extractable-protein potential rather than evidence of established suitability as a protein feedstock.

**Figure 3.**
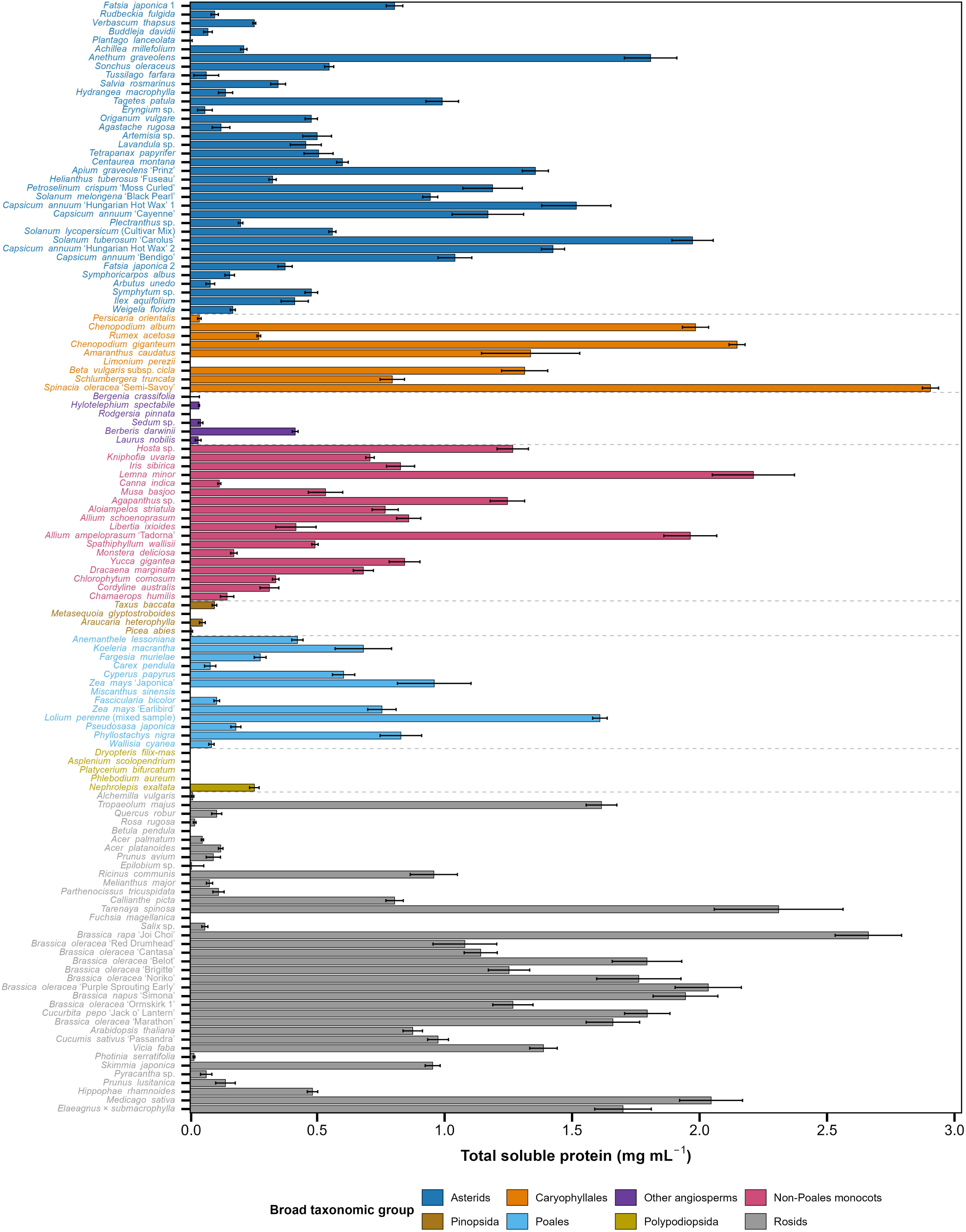
Total soluble protein concentrations across the plant biomass panel. Bars show mean total soluble protein concentrations from three replicates (n = 3), with error bars representing standard deviation. Samples are grouped by broad taxonomic group, indicated by colour and separated by dashed lines. TSP was quantified using the Bradford assay and is expressed as mg mL□¹.

A notable contrast was observed within *Solanum*: *S. tuberosum* ‘Carolus’ produced 1.973 mg mL□¹ TSP, approximately 3.5-fold more than *S. lycopersicum*. This difference shows that close taxonomic relatedness did not translate into similar protein recovery under the common workflow. Nevertheless, the lower tomato value does not necessarily indicate lower total leaf-protein content. During tomato-leaf processing in other studies, substantial losses have been attributed to retention of protein-containing liquid within fibrous pulp and the presence of insoluble protein, while younger upper leaves produced higher protein yields than older leaves (Yu et al., 2024). Reduced tomato leaf extractability during development has also been associated with decreased protein synthesis, increased protein degradation and incomplete extraction of the soluble-protein fraction (Kleuter et al., 2025). Thus, the relatively low TSP values observed for tomato and other biomass sources should not necessarily be interpreted as indicative of low protein abundance but may instead result from developmental changes and matrix-related limitations that reduce protein extractability. The constant biomass-to-extractant ratio (30 µL mg□¹ dry biomass) excluded biomass loading as an explanation for the observed variation. Nevertheless, a common protocol is unlikely to recover protein equally from structurally and chemically contrasting biomass matrices. Measured TSP may reflect the intrinsic soluble protein pool, developmental stage, protein degradation or aggregation, tissue accessibility, and interactions with phenolic compounds, pigments and cell wall polysaccharides. In plant-based protein, increased alkaline protein recovery was associated principally with disruption of the tissue and cell wall matrix rather than increased protein solubility or hydrolysis (Zhang et al., 2015). Different physical treatments similarly produced different soluble protein and Rubisco recoveries from sugar-beet leaves (Dukić et al., 2023), confirming that recovery is process dependent.

TSP should therefore be interpreted as an operational measure of soluble protein recovered under the applied conditions, rather than a direct measure of total tissue protein, extraction yield, protein purity or Rubisco abundance. Rubisco may account for up to approximately 50 % of soluble leaf protein in some plants (Parry et al., 2012), this proportion varies substantially with species, tissue type and developmental stage, as demonstrated by changes in the association between soluble protein and Rubisco abundance during rice development (Ishimaru et al., 2001).

Despite the reliance on Bradfords to measure TSP it fails to distinguish individual proteins, and as such caution is required when interpreting TSP data. With MAPP, the individual protein is assessed, however it is not quantifiable, so efforts were made to understand the relationship between MAPP and TSP data. **Fig. 3** establishes contrasting extractable protein phenotypes, whereas **Fig. 4** tests whether this independent biochemical measurement is associated with the Rubisco directed MAPP signals. All three Rubisco directed MAPP signals were strongly and positively correlated with TSP (*r* = 0.69–0.78, *p* < 0.001) (**Fig. 4**), indicating that biomasses with greater Bradford reactive soluble protein recovery generally produced stronger Rubisco directed MAPP signals. This agreement is biologically consistent with Rubisco comprising a substantial fraction of the soluble leaf protein pool, while the consistent relationships across two RbcL antibodies and RbcS suggest that the pattern was not specific to a single antibody. However, because TSP includes non-Rubisco proteins and MAPP signals are relative, the two measurements should be considered complementary rather than quantitatively interchangeable.

**Figure 4.**
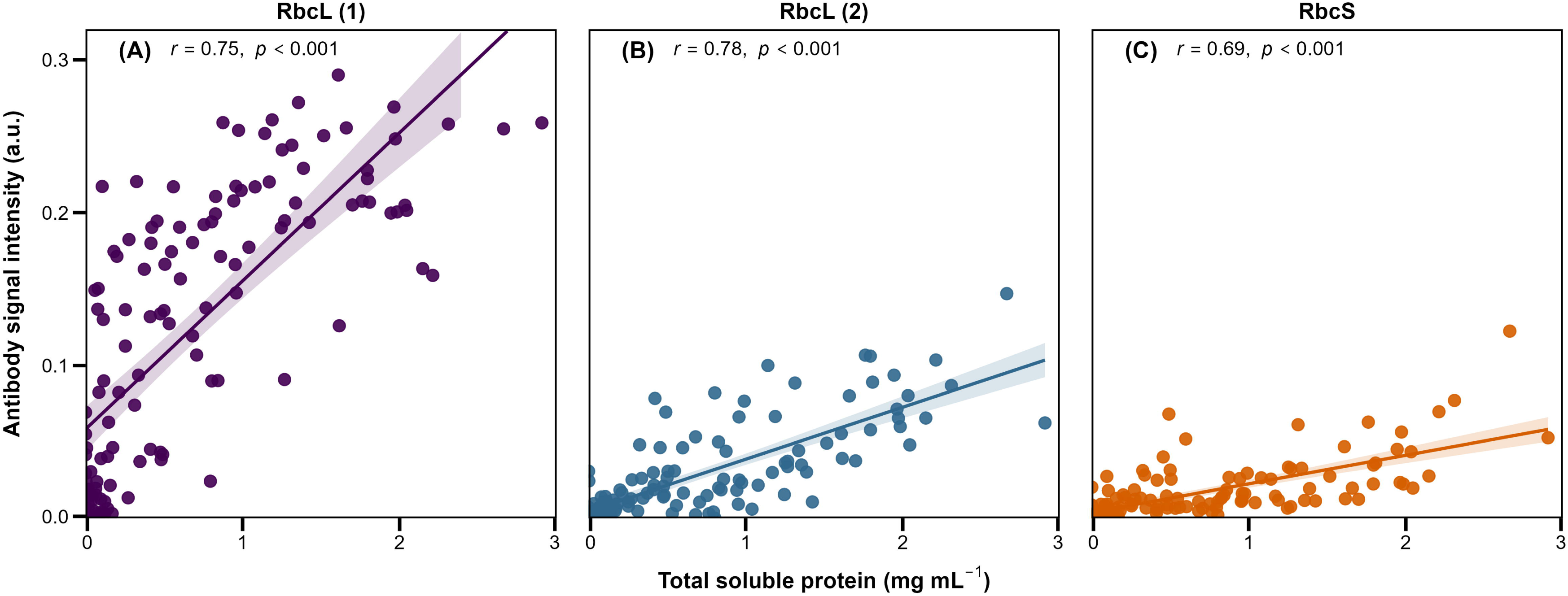
Relationships between Rubisco antibody signal intensity and total soluble protein across the biomass panel. Scatterplots show mean total soluble protein (TSP) concentration against mean signal intensity for (A) RbcL (1), (B) RbcL (2) and (C) RbcS. Each point represents the mean of three biological replicates for one biomass sample. Solid lines show fitted linear regressions and shaded bands represent 95% confidence intervals. Pearson correlation coefficients (*r*) and corresponding *p* values are shown. Antibody designations: RbcL (1), Abcam anti-Rubisco large subunit; RbcL (2), Agrisera anti-Rubisco large subunit; RbcS, Agrisera anti-Rubisco small subunit. Further antibody details are provided in Supplementary table 2.

### 3.4 MAPP analysis of cell wall polysaccharides

Cell walls are comprised of complex three-dimensional polysaccharide matrixes that collectively constitute most of the plant body. In some cases, cell wall polymers are known to impact protein accessibility and yield from biomass, and our goal was to investigate these effects for Rubisco. We therefore used MAPP to assess the relative abundance of cell wall structures across the plant sample set (**Fig. 5**) and integrated these data with protein (including Rubisco) yield data (**Data S1**).

**Figure 5.**
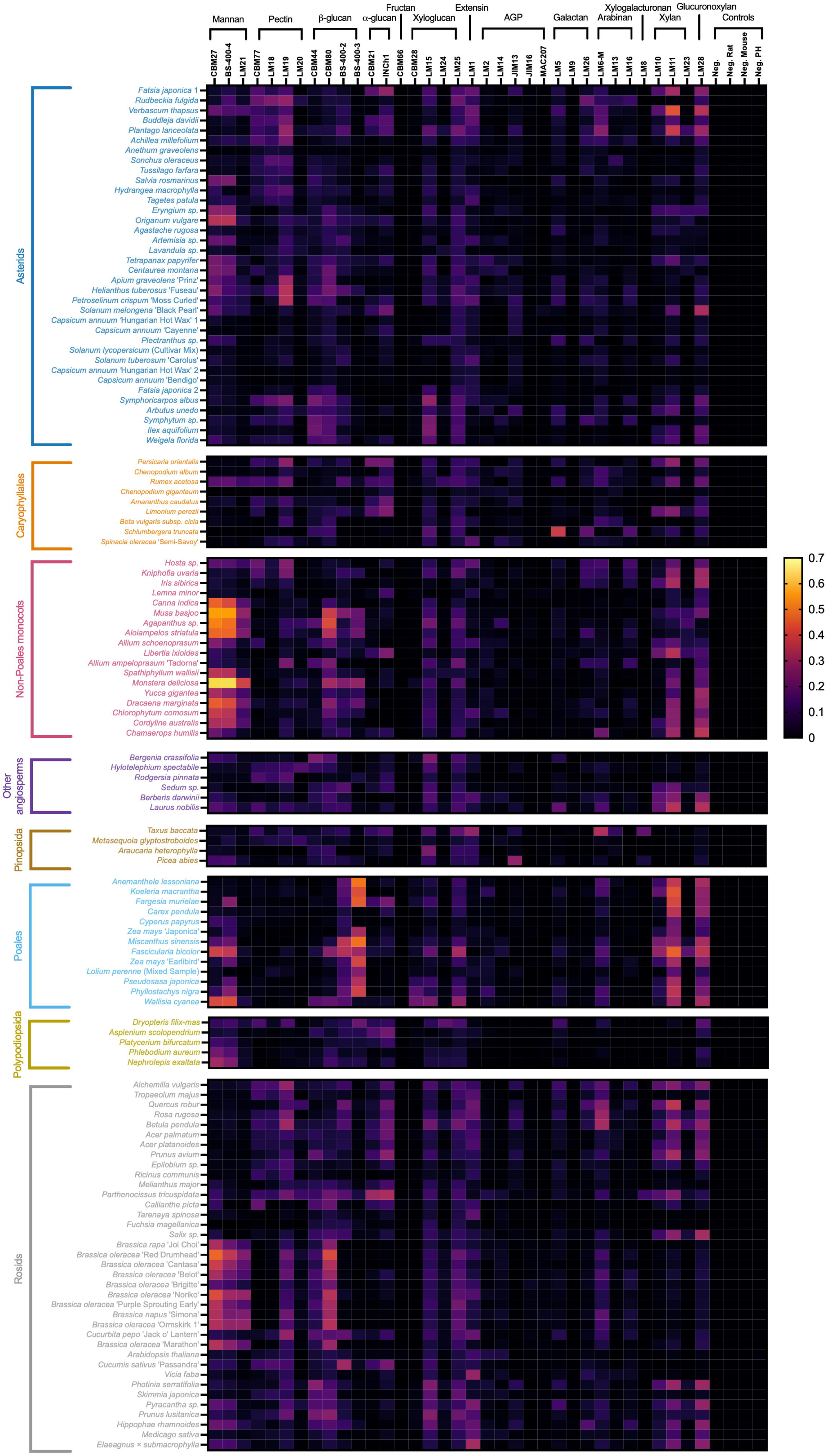
Glycan antibody binding profiles across the biomass panel. Rows represent biomass samples and columns represent glycan-binding antibodies grouped by their principal targets; relative signal intensity (0–0.8) increases from black through pink to yellow. Controls comprised a general negative control (Neg.) and detection-specific negative controls for rat, mouse and polyhistidine antibodies (Neg. Rat, Neg. Mouse and Neg. poly-His, respectively). Antibody identities, epitope specificities, dilutions, suppliers and references are provided in Supplementary table 2.

Summed signals for polysaccharide-specific probes across the CDTA, NaOH and cellulase extraction fractions revealed extensive variation in cell wall epitopes. As expected, (1→3,1→4)-β-glucan signals (detected by mAb BS-400-3) was highest in Poales with the nine highest signals originating from this group, led by *Anemanthele lessoniana* and *Miscanthus sinensis*. These data agree with the well-established occurrence of (1→3,1→4)-β-glucan in the cell walls of grasses and cereals. However, immunochemical and phylogenetic work has demonstrated mixed linkage glucan beyond Poaceae and even beyond Poales, so the non-Poales signals should be treated as biologically plausible rather than anomalous (Little et al., 2018; Sørensen et al., 2008).

Poales also showed high signals for the xylan directed mAb, LM11 and LM28, signals, with *Fascicularia bicolor* producing the maximum for both probes. The LM11/LM28 xylan signals are consistent with xylan being the dominant non-cellulosic polysaccharide in many grass secondary walls, and LM11 recognises both unsubstituted xylan and wheat arabinoxylan epitopes (McCartney et al., 2005; Tryfona et al., 2023). The strong Fascicularia, Koeleria, Fargesia, Anemanthele and bamboo signals support a recurring grass wall phenotype, while high signals in selected asterids, rosids and other monocots highlight the cross-lineage occurrence of xylan.

Non-Poales monocots showed a distinct profile for mannan containing polysaccharides (as detected by CBM27, and mAbs BS-400-4 and LM21). *Monstera deliciosa* produced the highest signal for all three probes, and *Musa basjoo*, *Agapanthus* sp. and *Aloiampelos striatula* also ranked highly. The mannan directed profile in non-Poales monocots is corroborated by previous MAPP data showing prominent soluble mannans in several non-commelinid monocot leaves and succulent tissues (Fradera-Soler et al., 2022). The agreement of CBM27, BS-400-4 and LM21, together with the repeated prominence of Monstera, Musa, Agapanthus and Aloiampelos, makes this more persuasive than a single probe maximum. Strong mannan signals in selected Brassica samples remain an important exception.

The glycan screen provides insight into the three-dimensional cell wall architectures from which soluble, and organellar proteins must be released, but it should not be presented as a direct assay of recalcitrance. Cell walls are dynamic composite materials whose porosity, hydration, charge and mechanics emerge from interactions among cellulose, hemicelluloses, pectin, proteins and phenolics (Anderson & Kieber, 2020; Thompson & Islam, 2021). A highly accessible epitope signal can therefore indicate composition, extractability or exposure; it does not by itself show that the epitope physically traps Rubisco. In green-tea leaf residue, alkaline protein recovery tracked tissue disruption and co-extraction of rhamnogalacturonan-I pectin and lignin rather than simple protein solubility, although that mechanism was specific to the material and treatment studied (Zhang et al., 2015). Work on tomato leaf likewise suggests that developmental changes in the soluble proteome and inhibitory compounds affect recovery (Kleuter et al., 2025), while recent studies on sugar-beet show that protein content and extractability can be screened at scale and have exploitable genetic variation (Rijken et al., 2025; Rijken et al., 2026).

Taken together, these results shed new light on the impact of cell wall architectures and glycan diversity in protein, including Rubisco, extraction from plant biomass, and highlight the need for detailed understanding of these effects to optimise extraction protocols.

### 3.5 Deciphering relationships between protein and glycan profiles and implications for Rubisco yield

To better understand correlative, and possibly causal, relationships between profiles obtained for TSP, cell wall glycans and Rubisco, we undertook Pearson correlation coefficient and Principal component analysis (PCA).

Pearson correlations were calculated across 128 plant biomass samples for all combinations of 13 protein antibodies and 15 glycan antibodies, yielding 195 pairwise correlations. Rows and columns were clustered using Euclidean distance and complete linkage applied to their full correlation profiles (**Fig. 6**). Proteins clustering together therefore showed similar patterns of correlation across the 15 glycans, while glycans clustering together showed similar relationships across the 13 protein antibodies. Of the 195 correlations, 146 were negative and 49 were positive, with a mean absolute correlation coefficient (|*r*|) of 0.139.

**Figure 6.**
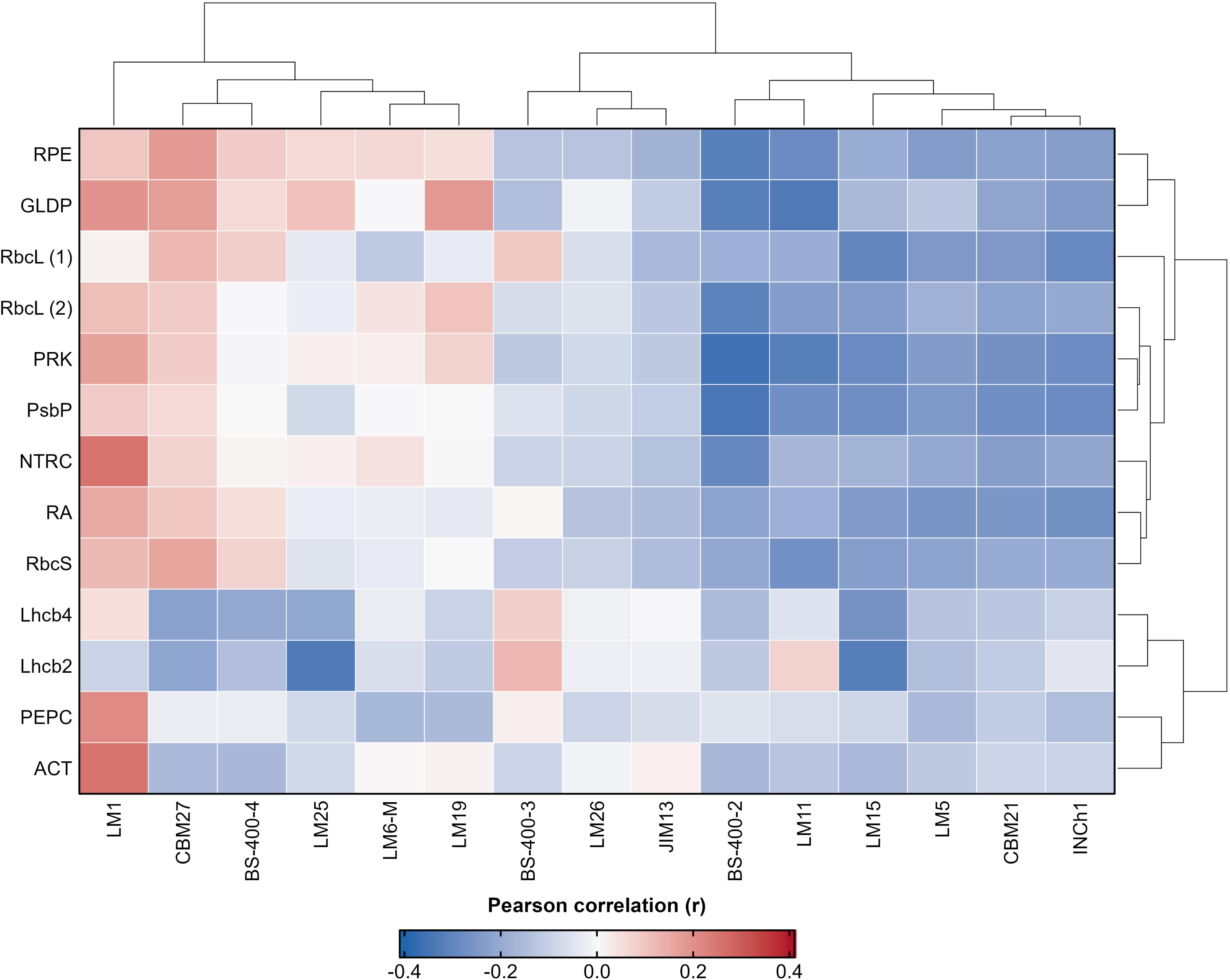
Pearson correlation heatmap of protein and glycan antibody signals across the plant biomass panel. Each cell represents the Pearson correlation coefficient (*r*) calculated between mean protein and glycan antibody signal intensities across 128 plant samples. Red indicates positive correlations, with protein and glycan antibody signals tending to increase together; blue indicates negative correlations, with higher protein antibody signals associated with lower glycan antibody signals. Near-white cells indicate little or no linear association. Protein and glycan antibodies were hierarchically clustered using Euclidean distance and complete linkage according to the similarity of their complete correlation profiles. RbcL (1) and RbcL (2) are Abcam and Agrisera antibodies against the Rubisco large subunit, respectively; RbcS is an Agrisera antibody against the Rubisco small subunit. Further antibody identities and specificities are provided in Supplementary table 2.

The glycan dendrogram separated a region of mixed or positive correlations, containing LM1 and several antibodies with weak or near null associations, from a predominantly negative block comprising mAbs LM15, LM5, INCh1, BS-400-2, LM11 and CBM21. Many photosynthetic proteins shared broadly similar negative correlation profiles, whereas Lhcb2/Lhcb4 and PEPC/ACT formed more distinct branches.

Following false discovery rate (FDR) correction, 41 correlations remained significant, comprising 39 negative and 2 positive associations. The strongest negative correlations involved PRK–BS-400-2 (*r* = −0.365), PsbP–BS-400-2 (*r* = −0.347), GLDP–LM11 (*r* = −0.340) and Lhcb2–LM25 (*r* = −0.337). ACT–LM1 (*r* = 0.258) and NTRC–LM1 (*r* = 0.256) were the only significant positive correlations. Nine Rubisco–glycan correlations remained significant, all negative. The strongest involved RbcL (2)–BS-400-2 (*r* = −0.316), RbcL (1)–LM15 (*r* = −0.304), RbcL (1)–INCh1 (*r* = −0.297) and RbcS–LM11 (*r* = −0.275). LM15 showed significant negative correlations with all three Rubisco antibodies, while LM11 and INCh1 showed consistently negative associations across the same antibodies. This agreement suggests that the observed patterns were not specific to a single Rubisco antibody.

Across the biomass samples, stronger Rubisco and photosynthetic-protein signals tended to coincide with weaker signals for certain structural glycan epitopes. However, these associations were generally weak and do not establish that individual glycans directly restrict Rubisco extraction. The predominantly negative correlation block may instead reflect differences in tissue composition or biomass type. LM1 showed a contrasting correlation profile, while several other glycan antibodies had weak or near-null associations, indicating that the relationships between protein and glycan signals were not uniformly negative. In a sensitivity analysis including TSP (**Fig. S1),** five TSP–glycan correlations remained significant after FDR correction. The strongest negative associations involved BS-400-2 (*r* = −0.404) and LM11 (*r* = −0.354), suggesting that similar relationships may extend beyond Rubisco to soluble protein more generally. These findings identify glycan epitopes that warrant further investigation as potential markers of biomass extractability, particularly through validation against protein recovery and responses to enzyme treatment. Their associations with protein signals do not, however, establish a causal role in limiting extraction.

PCA of the targeted protein-glycan dataset identified two complementary axes of variation across the biomass panel (**Fig. 7**). In the targeted PCA (**Fig. 7A**), PC1 explained 29.7% of the variance and was dominated by RbcL (1), RbcL (2) and RbcS, which together contributed 82.5% of the axis, whereas the glycan antibodies BS-400-4, LM1, LM11, LM15, LM19 and JIM13 contributed 98.2% of PC2, which explained 19.8% of the variance. Together, PC1 and PC2 accounted for 49.4% of the total variation. All three Rubisco antibodies loaded in the same direction on PC1, whereas LM11 and LM15 loaded oppositely, consistent with the inverse Rubisco-glycan relationships identified in **Fig. 6**. LM1, LM19, JIM13 and BS-400-4 together contributed 96.4% of PC2, indicating a contrast among glycan epitopes rather than a general increase in glycan antibody signal. The six targeted glycans collectively explained 15.9% of the multivariate Rubisco antibody variation in redundancy analysis (adjusted R² = 0.117, BH-adjusted P < 0.001), with LM15 retaining the strongest marginal association (BH-adjusted P = 0.0066). In the HC3 robust multivariable model, LM15 was negatively associated with Rubisco PC1 (standardised β = -0.300, 95% CI -0.432 to -0.168, BH-adjusted P < 0.001). This relationship was also supported by Pearson and Spearman analyses, identifying LM15 as the most consistent candidate marker of lower Rubisco associated signal across the different analytical approaches.

**Figure 7.**
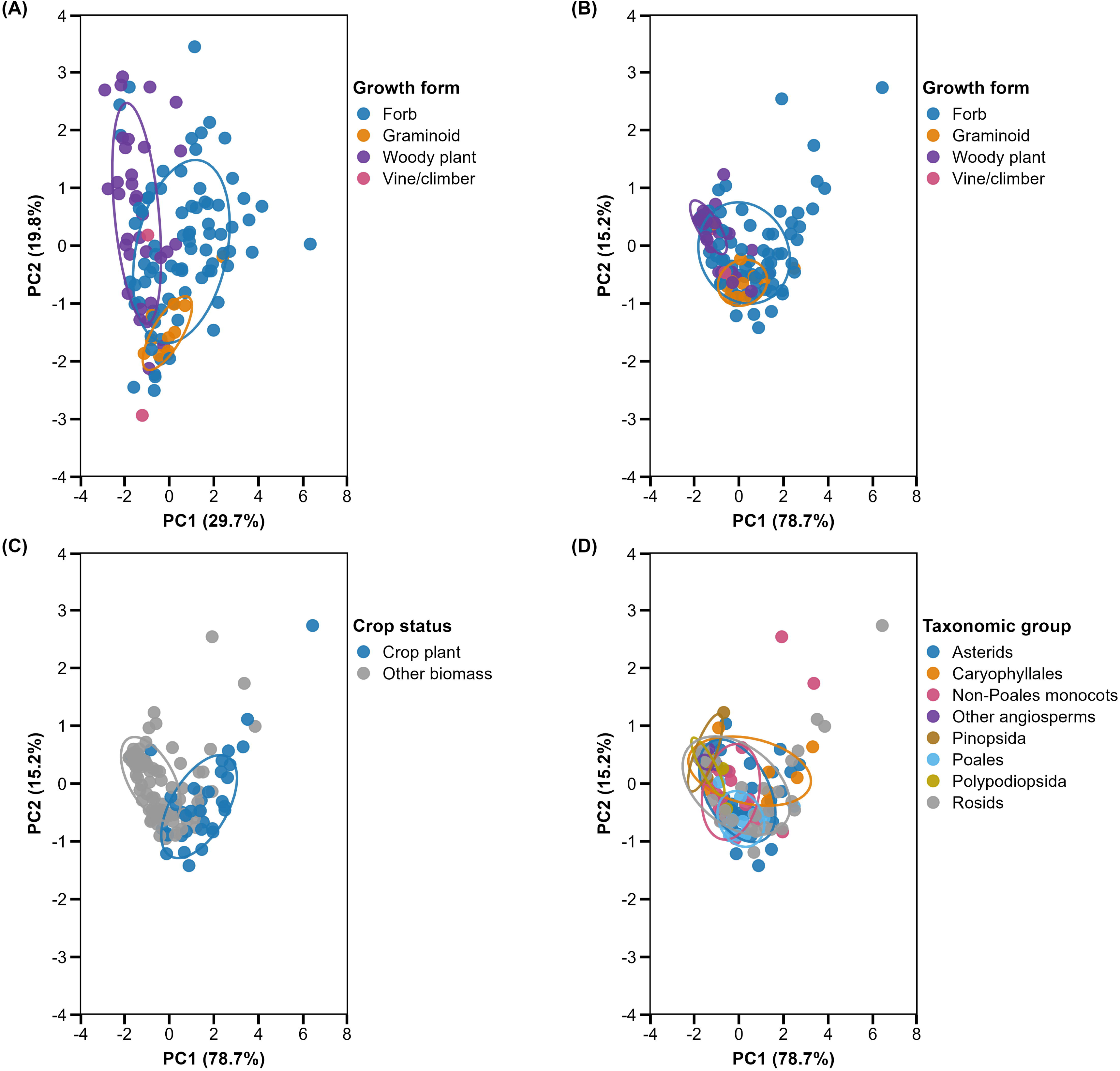
Principal component analysis of targeted protein-glycan and Rubisco antibody profiles across the plant biomass panel. (A) Targeted PCA based on RbcL (1), RbcL (2), RbcS and six selected glycan antibodies (BS-400-4, LM1, LM11, LM15, LM19 and JIM13), points are coloured by growth form. (B) Rubisco-only PCA based on the same three Rubisco antibodies, points are coloured by growth form. The same Rubisco-only scores are displayed by (C) crop status and (D) broad taxonomic group. Each point represents one plant sample (n = 128), based on the mean of three biological replicates; ellipses represent 68% within-group dispersion. In PCA analyses, positive PC1 scores indicate increasing combined Rubisco antibody signal, whereas PC2 represents the largest remaining source of variation independent of PC1. Percentages in the axis labels indicate the variance explained by each principal component. Samples positioned closer together have more similar antibody profiles, whereas greater separation indicates greater dissimilarity. Biomass identities and classifications are provided in Table 1.

When grouped by growth form (**Data S2**), forbs (non-graminoid herbaceous flowering plants) were shifted towards the Rubisco associated zone of PC1, woody plants towards the opposing end, and graminoids towards negative PC2 values. A separate Rubisco-only PCA explained 78.7% and 15.2% of the variance on PC1 and PC2, respectively; together, these axes accounted for 93.9% of the variance (**Fig. 7B-D**). In this space, forbs and crop plants were descriptively shifted towards the Rubisco-associated end of PC1, whereas woody plants were shifted towards the opposing end; broad taxonomic groups overlapped extensively. Growth form explained 14.6% of the targeted profile variation and 16.1% of the Rubisco only variation (PERMANOVA, both BH-adjusted P < 0.001). However, multivariate dispersion also differed among growth forms (both PERMDISP P = 0.001), indicating that these effects reflected differences in both group position and within group heterogeneity. Crop status produced the strongest group effect on the Rubisco only profile (R² = 0.249, BH-adjusted P < 0.001), without a supported difference in dispersion (P = 0.064). Substantial within group variation remained, particularly among the forbs.

These patterns suggest that variation in Rubisco associated signal is not organised by broad taxonomy alone. Crop status and growth form may provide useful, although imperfect, guidance for candidate selection, while the targeted PCA shows that Rubisco associated signal and cell-wall epitope profiles represent complementary dimensions of biomass phenotype. Complementary HC3 modelling showed that LM11 and LM15 were negatively associated with TSP, whereas LM1 was positively associated; all three directions were also supported by Pearson and Spearman analyses. Together with **Fig. 6**, these findings identify LM15 as the most consistent Rubisco associated marker, LM11 as a candidate marker of the broader soluble protein phenotype, and LM1 as a marker associated with higher TSP rather than Rubisco specifically. The separate protein and glycan heatmaps (**Fig. 2 and Fig. 5**) showed substantial sample level variation, while **Fig. 4**, **Fig. 6 and Fig. 7** show that TSP, Rubisco antibody and glycan antibody signals covary through complementary pairwise and multivariate analyses. This interpretation is consistent with reports that biomass structure is associated with the success of leaf protein fractionation (Nynäs et al., 2021) and that pectin composition is associated with protein extractability in tomato leaves (Kleuter et al., 2024). However, these associations do not establish that cell-wall composition directly restricts Rubisco extraction. Crop derived biomass may provide a practical starting point for screening, but individual feedstocks must still be evaluated directly and may require biomass-specific processing strategies. These findings provide a basis for selecting contrasting feedstocks and evaluating whether cell-wall degrading enzyme treatments can improve protein recovery biomasses with contrasting cell-wall epitope profiles.

### 3.6 Impact of enzyme treatment on soluble protein yield and Rubisco microarray signals

It has previously been highlighted that cell wall degrading enzymes can enhance protein yield by disassembling the polysaccharide matrix that is the dominant component of biomass, and from which proteins must be released to be extracted effectively (Harholt et al., 2010; Krishnan et al., 2026; Pérez-Vila et al., 2022; Scheller & Ulvskov, 2010; Tamayo Tenorio et al., 2016). To assess these effects in relation to Rubisco yield, we used 7 different enzyme treatments tested against 3 different biomass types (**Table S3**). The three biomass types tested were grass, cucumber and tomato due to the abundance of waste generated annually from these crops. It was found that enzyme pretreatment significantly increased soluble protein yield in 51 of 84 biomass matrix preparation comparisons (**Fig. 8 and Fig. S2**).

**Figure 8.**
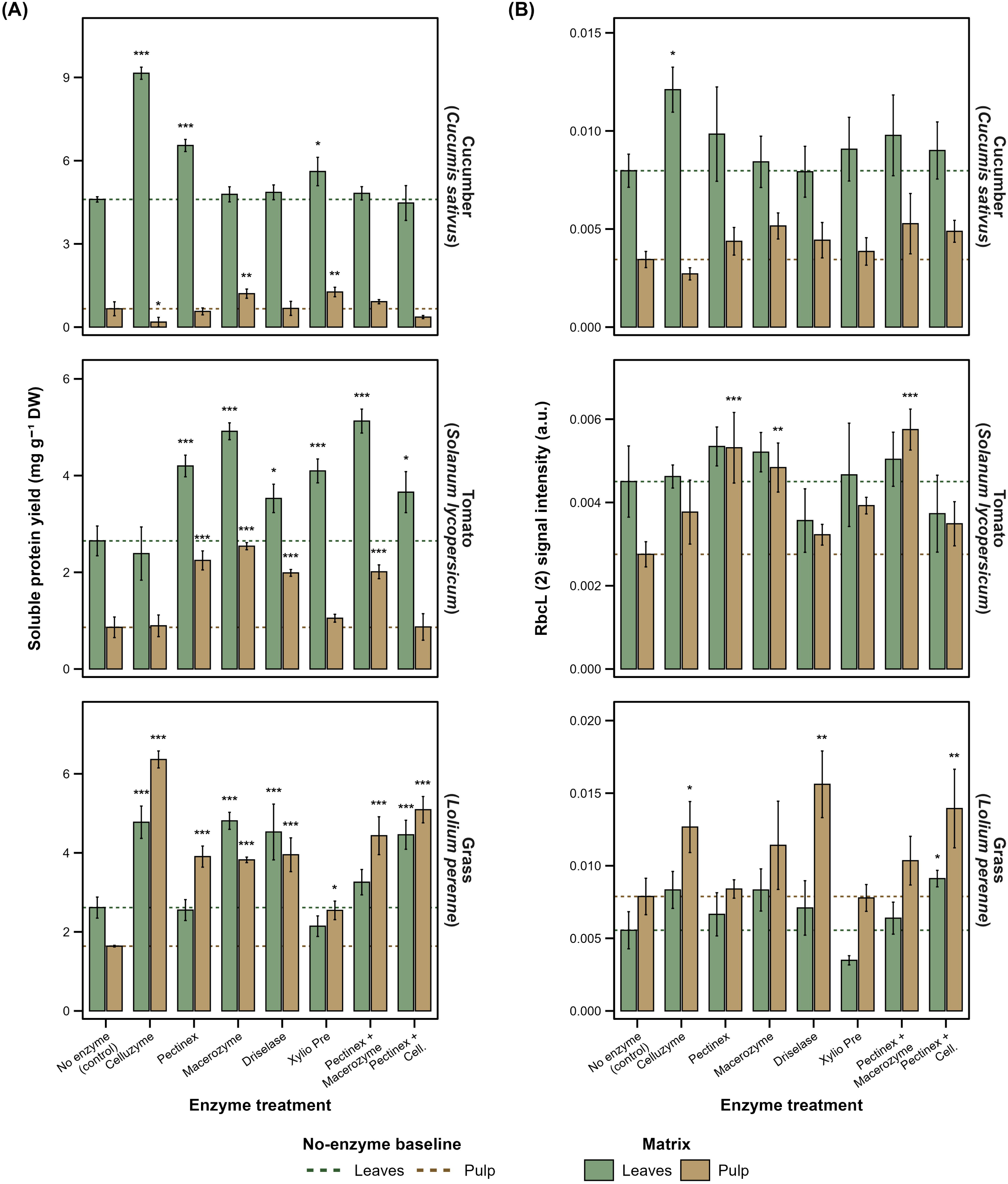
Effects of enzyme treatment on total soluble protein yield and RbcL (2) signal in frozen-thawed plant biomass. (A) Total soluble protein yield and (B) MAPP-derived RbcL (2) signal intensity in leaf and pulp fractions of cucumber (*Cucumis sativus*), tomato (*Solanum lycopersicum*) and perennial ryegrass (*Lolium perenne*). Bars show means ± SD from three biological replicates (n = 3). Green and brown bars represent leaf and pulp fractions, respectively; dotted horizontal lines indicate the corresponding mean no-enzyme controls. Asterisks indicate Dunnett-adjusted differences from the matched species-and matrix-specific no-enzyme control: *P* < 0.05, **P** < 0.01 and ***P*** < 0.001. RbcL (2) denotes the Agrisera antibody targeting the Rubisco large subunit. Biomass weights and enzyme concentrations are provided in Supplementary table 3. DW, dry weight.

Grass demonstrated the strongest response to enzymatic pre-treatment of all matrices examined, with Celluzyme treatment producing the greatest increase in both in TSP and RbcL (2) abundance (**Fig. 8 and Fig. S2**). The convergence of these independent measurements provides strong evidence that enzymatic degradation of the grass cell wall enhanced the recovery of intracellular proteins, particularly chloroplast associated Rubisco. Similar responses were also observed in grass pulp, suggesting that enzyme assisted extraction is effective not only for intact leaf tissue but also for residual biomass streams that currently have limited commercial value. The marked improvement in protein recovery is consistent with the structural characteristics of grass cell walls. Grasses possess Type II primary cell walls enriched in glucuronoarabinoxylans and mixed-linkage β-glucans, with comparatively low pectin content and frequent ferulate mediated cross linking between polysaccharides and lignin (Carpita & Gibeaut, 1993; Vogel, 2008). These features reduce wall porosity and limit solvent penetration, making mechanical homogenisation alone insufficient to fully release intracellular proteins. Cellulolytic enzyme preparations such as Celluzyme likely weaken this structural network, increasing wall permeability and facilitating disruption of chloroplast containing mesophyll cells. Similar mechanisms have been reported during enzyme assisted extraction of proteins from plant biomass, where degradation of structural polysaccharides enhances diffusion of soluble proteins into the extraction medium (Sari et al., 2013; Wang et al., 2025). Previous studies have demonstrated that matching enzyme composition to cell wall architecture significantly improves biomass deconstruction and intracellular product recovery (Park et al., 2016; Shawky et al., 2025). Consequently, optimisation of enzyme formulations for monocotyledonous biomass may further enhance protein yields while reducing enzyme loading and processing costs.

Cucumber exhibited a positive but less consistent response to enzymatic pre-treatment than grass. Celluzyme produced the highest TSP recovery from frozen-thawed leaves (**Fig. 8A**), suggesting that combining freeze-thaw disruption with enzymatic hydrolysis improved access to intracellular proteins. Freeze-thaw processing is known to induce membrane rupture through ice crystal formation, thereby increasing tissue permeability and enhancing subsequent enzyme penetration (Mazur, 1984). The present findings therefore suggest that physical disruption and enzymatic degradation act synergistically to improve protein extraction from cucumber tissues.

In contrast to TSP recovery, RbcL abundance did not consistently increase following the same treatments. Macerozyme generated the highest Rubisco signal in freeze-dried leaves and pulp, whereas Celluzyme produced the strongest response in frozen-thawed leaves (**Fig. 8B and Fig. S2B**). This divergence suggests that the mechanisms governing total protein extraction differ from those controlling chloroplast protein recovery. While Celluzyme may effectively disrupt structural cellulose networks and release soluble cytoplasmic proteins, Macerozyme may more effectively degrade pectic components surrounding chloroplast rich tissues or better preserve Rubisco during extraction. Similar differences between total protein recovery and individual protein abundance have been reported in enzyme assisted extraction studies, reflecting the complexity of plant tissue organisation and the compartmentalisation of intracellular proteins (Komatsu & Jorrin-Novo, 2021; Wang et al., 2025). Compared with grass pulp, cucumber pulp showed smaller improvements in both TSP and Rubisco abundance. This may reflect differences in tissue composition, as cucumber tissues possess relatively thin primary cell walls, high water content and lower structural polysaccharide concentrations than grass (Cosgrove, 2005). Consequently, a larger proportion of proteins may already be accessible following mechanical disruption, reducing the additional benefit provided by enzymatic hydrolysis. Alternatively, endogenous protease activity released during tissue disruption may contribute to partial degradation of soluble proteins before extraction. Protease mediated losses have previously been identified as a significant limitation during protein recovery from fresh green biomass (Bals & Dale, 2012).

Tomato displayed a clear dependence on both processing method and enzyme formulation. Enzymatic effects were substantially greater in frozen-thawed samples than in freeze-dried material, with Macerozyme and the Pectinex + Macerozyme combination consistently producing the highest TSP recoveries from both leaf and pulp tissues (**Fig. 8A and Figure S2A**). These findings indicate that freeze-thaw processing enhanced enzyme accessibility and that degradation of pectin-rich cell wall domains was particularly effective in improving protein extraction. The observed response is consistent with the biochemical composition of tomato cell walls, which are characteristic of Type I primary walls containing abundant pectin, xyloglucans and cellulose microfibrils (Carpita & Gibeaut, 1993). Pectin contributes significantly to cell adhesion through the middle lamella, and enzymatic degradation of these polymers reduces tissue cohesion, facilitating cell separation and intracellular protein release (Willats et al., 2001). The superior performance of Pectinex-containing treatments therefore supports the hypothesis that pectin degradation is a key mechanism governing protein extraction from tomato tissues. This interpretation is further supported by the strongest RbcL (2) responses observed in tomato pulp following treatment with Pectinex + Macerozyme (**Fig. 8B and Fig. S2B**). Tomato processing generates substantial quantities of pomace and pulp that retain appreciable amounts of protein despite being predominantly utilised for low value applications such as animal feed or compost (Reguengo et al., 2022). Improving protein recovery from these residues could therefore contribute to increased resource efficiency within tomato processing industries and support circular bioeconomy strategies. Similar approaches have recently been proposed for the valorisation of vegetable processing side streams through enzyme assisted extraction technologies (Shawky et al., 2025; Wang et al., 2025). Collectively, the tomato results demonstrate that enzyme specificity should be matched to tissue composition, with pectin degrading enzymes offering particular advantages for dicotyledonous tissues rich in pectic polysaccharides. The strong responses observed in tomato pulp further suggest that vegetable processing residues represent promising substrates for sustainable protein recovery.

Overall, these findings demonstrate the potential of enzyme assisted extraction to improve protein recovery across different plant biomass types, although extraction efficiency and Rubisco recovery depend on substrate composition, enzyme specificity and the intracellular localisation of target proteins. Grass exhibited the strongest response, particularly following Celluzyme treatment, suggesting that cellulose rich structural components represent an important barrier to protein release. Cucumber also benefited from enzymatic treatment, although the optimal enzyme preparation may differ depending on whether maximising total protein recovery or preserving Rubisco is the primary objective. In tomato, increased TSP recovery was not always accompanied by corresponding increases in RbcL abundance, highlighting the importance of distinguishing total protein recovery from target protein abundance. Such discrepancies may reflect biological variation or matrix-dependent effects on immunodetection rather than changes in Rubisco recovery alone (Komatsu & Jorrin-Novo, 2021). Consequently, further validation using complementary analytical techniques, such as SDS-PAGE or targeted LC-MS/MS, would help clarify treatment-specific changes in Rubisco abundance. Future optimisation should therefore consider both enzyme specificity and substrate composition, alongside the preservation and reliable quantification of target proteins, to maximise the potential of enzyme assisted extraction for sustainable protein recovery from agricultural biomass.

## 4. Conclusion

There is broad agreement that Rubisco is a promising source of plant protein that could make a valuable contribution to balancing future global nutritional needs with environmental considerations. However, it is also clear that significant optimisation of extraction protocols is required if this potential is to be realised in a cost-feasible way. In this work we demonstrate how a high-throughput technology platform (MAPP), can provide important new insights into three parameters that play key roles in determining Rubisco yield: biomass source; cell wall matrix composition and enzymatic disruption of cell wall polymers.

Our results show that Rubisco abundance varies considerably across plant biomass types, and that abundance is not directly linked to taxonomic groupings. To our knowledge, this is the first parallel analysis of cell wall matrix polysaccharides and soluble proteins across a diverse plant biomass panel, allowing relationships between these glycans and Rubisco yield to be investigated. We explored these effects further by demonstrating how the targeted use of cell wall degrading enzymes can enhance Rubisco yield. Additionally, crop plants generally produced stronger Rubisco-associated signals, with potato haulm and Brassica leaf residues emerging as promising feedstocks for further extraction and valorisation.

This work highlights the importance of understanding biomass properties when developing Rubisco extraction methods and demonstrates that MAPP can aid in addressing this knowledge gap. We anticipate that MAPP could also be used to examine other sources of variation in Rubisco yield and quality, including seasonal and developmental variation, batch-to-batch variation, and Rubisco purity and composition.

## 5. CRediT authorship contribution statement

**John C. D. Pattinson:** Conceptualisation, data curation, formal analysis, investigation, methodology, project administration, resources, software, validation, visualization, writing – original draft, writing – review and editing. **Charlotte S. Fletcher:** Conceptualisation, data curation, formal analysis, investigation, methodology, project administration, resources, software, validation, visualization, writing – original draft, writing – review and editing. **Ieva Lelenaite:** Project administration, writing – review and editing. **Georg Lietz:** Supervision **William G. T. Willats:** Funding acquisition, Supervision, writing – review and editing. **Maxim. V. Kapralov:** Funding acquisition, Supervision, writing – review and editing.

## 7. Acknowledgements

The authors would like to acknowledge Ewelina Bien and Max Kelly for their support and guidance. This research was supported by a PhD studentship funded through the Institute for Agri-Food Research and Innovation (IAFRI) Doctoral Training Programme at Newcastle University (JCDP). This work was supported by a PhD studentship funded by the BBSRC in relation to the BiSCoP (Bioscience for Sustainable Consumer Products) Collaborative Training Partnership.

## 8. Appendix A. Supplementary data

Data S1

Data S2

## 9. Appendix B. Supplementary figures and tables

Table S1

Table S2

Table S3

Fig. S1

Fig. S2

## 10. Data availability

Data will be made available upon request.

