## Supplementary figures and tables for "High-throughput microarray analysis provides new insight into optimising Rubisco yield from diverse plants"

**Supplementary Table Captions**

**Table S1.** Complete list of species used for this experiment.

**Table S2.** Complete list of primary and secondary antibodies used, the epitopes they bind, and the studies in which they were described

**Table S3.** Plant material weights required with enzyme concentrations

**Supplementary Figure Captions**

**Fig. S1.** Sensitivity analysis of Pearson correlations between total soluble protein, protein-probe and glycan-probe signals across the plant biomass panel.

**Fig. S2. Effects of enzyme treatment on total soluble protein yield and RbcL (2) signal in freeze-dried plant biomass.**

**Table S1.** Complete list of species used for this experiment.

| Family | Species | Common name |
| --- | --- | --- |
| Araliaceae | *Fatsia* japonica | Paperplant |
| Asteraceae | *Rudbeckia* fulgida | Black-eyed Susan |
| Scrophulariaceae | *Verbascum* thapsus | Common mullein |
| Scrophulariaceae | *Buddleja* davidii | Butterfly bush |
| Plantaginaceae | *Plantago* lanceolata | Narrow-leaved plantain |
| Asteraceae | *Achillea* millefolium | yarrow |
| Apiaceae | *Anethum* graveolens | Dill |
| Asteraceae | *Sonchus* oleraceus | Common Sow-Thistle |
| Asteraceae | *Tussilago* farfara | Coltsfoot |
| Lamiaceae | *Salvia* rosmarinus | Rosemary |
| Hydrangeaceae | *Hydrangea* macrophylla | Bigleaf Hydrangea |
| Asteraceae | *Tagetes* patula | French Marigold |
| Apiaceae | *Eryngium* sp. | Eryngo |
| Lamiaceae | *Origanum* vulgare | Oregano |
| Lamiaceae | *Agastache* rugosa | Korean mint |
| Asteraceae | *Artemisia* sp. | Wormwood |
| Lamiaceae | *Lavandula* sp. | Lavender |
| Araliaceae | *Tetrapanax* papyrifer | Rice Paper Plant |
| Asteraceae | *Centaurea* montana | Perennial Cornflower |
| Apiaceae | *Apium* graveolens | Celeriac |
| Asteraceae | *Helianthus* tuberosus | Jerusalem Artichoke |
| Apiaceae | *Petroselinum* crispum | Parsley |
| Solanaceae | *Solanum* melongena | Aubergine |
| Solanaceae | *Capsicum* annuum | Hungarian Hot Wax Pepper |
| Solanaceae | *Capsicum* annuum | Cayenne Pepper |
| Lamiaceae | *Plectranthus* sp. | Swedish Ivy |
| Solanaceae | *Solanum* lycopersicum | Tomato |
| Solanaceae | *Solanum* tuberosum | Potato |
| Solanaceae | *Capsicum* annuum | Hungarian Hot Wax Pepper |
| Solanaceae | *Capsicum* annuum | Bell pepper |
| Araliaceae | *Fatsia* japonica | Paperplant |
| Caprifoliaceae | *Symphoricarpos* albus | Common Snowberry |
| Ericaceae | *Arbutus* unedo | Strawberry Tree |
| Boraginaceae | *Symphytum* sp. | Comfrey |
| Aquifoliaceae | *Ilex* aquifolium | English Holly |
| Caprifoliaceae | *Weigela* florida | Weigela |
| Polygonaceae | *Persicaria* orientalis | Kiss-me-over-the-garden-gate |
| Amaranthaceae | *Chenopodium* album | Lamb's quarters |
| Polygonaceae | *Rumex* acetosa | Garden sorrel |
| Amaranthaceae | *Chenopodium* giganteum | Tree Spinach |
| Amaranthaceae | *Amaranthus* caudatus | Love-Lies-Bleeding |
| Plumbaginaceae | *Limonium* perezii | Sea Lavender |
| Chenopodiaceae | *Beta* vulgaris subsp. cicla | Swiss Chard |
| Cactaceae | *Schlumbergera* truncata | Thanksgiving Cactus |
| Chenopodiaceae | *Spinacia* oleracea | Spinach |
| Asparagaceae | *Hosta* sp. | Plantain lilies |
| Asphodelaceae | *Kniphofia* uvaria | Red hot poker |
| Iridaceae | *Iris* sibirica | Siberian Iris |
| Araceae | *Lemna* minor | Common Duckweed |
| Cannaceae | *Canna* indica | Indian shot |
| Musaceae | *Musa* basjoo | Japanese Banana |
| Amaryllidaceae | *Agapanthus* sp. | African Lily |
| Asphodelaceae | *Aloiampelos* striatula | Hardy Aloe |
| Amaryllidaceae | *Allium* schoenoprasum | Chive |
| Iridaceae | *Libertia* ixioides | New Zealand Iris |
| Amaryllidaceae | *Allium* ampeloprasum | Leek |
| Araceae | *Spathiphyllum* wallisii | Peace Lily |
| Araceae | *Monstera* deliciosa | Swiss Cheese Plant |
| Asparagaceae | *Yucca* gigantea | Spineless Yucca |
| Asparagaceae | *Dracaena* marginata | Madagascar Dragon Tree |
| Asparagaceae | *Chlorophytum* comosum | Spider Plant |
| Asparagaceae | *Cordyline* australis | Cabbage Tree |
| Arecaceae | *Chamaerops* humilis | European Fan Palm |
| Crassulaceae | *Sedum* sp. | Stonecrop |
| Saxifragaceae | *Bergenia* crassifolia | Siberian tea |
| Crassulaceae | *Hylotelephium* spectabile | Showy stonecrop |
| Saxifragaceae | *Rodgersia* pinnata | Featherleaf Rodgersia |
| Berberidaceae | *Berberis* darwinii | Darwin's Barberry |
| Lauraceae | *Laurus* nobilis | Bay Laurel |
| Taxaceae | *Taxus* baccata | English yew |
| Cupressaceae | *Metasequoia* glyptostroboides | Dawn Redwood |
| Araucariaceae | *Araucaria* heterophylla | Norfolk Island pine |
| Pinaceae | *Picea* abies | Norway Spruce |
| Poaceae | *Anemanthele* lessoniana | Pheasant's tail grass |
| Poaceae | *Koeleria* macrantha | Prairie Junegrass |
| Poaceae | *Fargesia* murielae | Umbrella bamboo |
| Cyperaceae | *Carex* pendula | Pendulous Sedge |
| Cyperaceae | *Cyperus* papyrus | Papyrus |
| Poaceae | *Zea* mays subsp. mays | Maize |
| Poaceae | *Miscanthus* sinensis | Chinese Silvergrass |
| Bromeliaceae | *Fascicularia* bicolor | Fascicularia |
| Poaceae | *Zea* mays subsp. mays | Sweetcorn |
| Poaceae | *Lolium* perenne | Grass mix (dominant ryegrass) |
| Poaceae | *Pseudosasa* japonica | Arrow Bamboo |
| Poaceae | *Phyllostachys* nigra | Black Bamboo |
| Bromeliaceae | *Wallisia* cyanea | Pink Quill |
| Dryopteridaceae | *Dryopteris* filix-mas | Male fern |
| Aspleniaceae | *Asplenium* scolopendrium | Hart's Tongue |
| Polypodiaceae | *Platycerium* bifurcatum | Staghorn Fern |
| Polypodiaceae | *Phlebodium* aureum | Blue Star Fern |
| Nephrolepidaceae | *Nephrolepis* exaltata | Boston Fern |
| Rosaceae | *Alchemilla* vulgaris | Lady’s mantle |
| Tropaeolaceae | *Tropaeolum* majus | Nasturtium |
| Fagaceae | *Quercus* robur | English oak |
| Rosaceae | *Rosa* rugosa | Rugosa rose |
| Betulaceae | *Betula* pendula | European white birch |
| Sapindaceae | *Acer* palmatum | Japanese maple |
| Sapindaceae | *Acer* platanoides | Norway maple |
| Rosaceae | *Prunus* avium | Sweet cherry |
| Onagraceae | *Epilobium* sp. | Willowherb |
| Euphorbiaceae | *Ricinus* communis | Castor Bean |
| Francoaceae | *Melianthus* major | Honeybush |
| Vitaceae | *Parthenocissus* tricuspidata | Boston Ivy |
| Malvaceae | *Callianthe* picta | Painted Abutilon |
| Cleomaceae | *Tarenaya* spinosa | Spider Flower |
| Onagraceae | *Fuchsia* magellanica | Hardy Fuchsia |
| Salicaceae | *Salix* sp. | Willow |
| Brassicaceae | *Brassica* rapa subsp. chinensis | Pak Choi |
| Brassicaceae | *Brassica* oleracea var. capitata | Red Cabbage |
| Brassicaceae | *Brassica* oleracea var. capitata | White Cabbage |
| Brassicaceae | *Brassica* oleracea var. botrytis | Cauliflower |
| Brassicaceae | *Brassica* oleracea var. gemmifera | Brussels Sprouts |
| Brassicaceae | *Brassica* oleracea var. gongylodes | Kohlrabi |
| Brassicaceae | *Brassica* oleracea var. italica | Sprouting Broccoli |
| Brassicaceae | *Brassica* napus subsp. rapifera | Swede |
| Brassicaceae | *Brassica* oleracea var. capitata | White Cabbage |
| Cucurbitaceae | *Cucurbita* pepo | Pumpkin |
| Brassicaceae | *Brassica* oleracea var. italica | Broccoli |
| Brassicaceae | *Arabidopsis* thaliana | Thale cress |
| Cucurbitaceae | *Cucumis* sativus | Cucumber |
| Fabaceae | *Vicia* faba | Faba Bean |
| Rosaceae | *Photinia* serratifolia | Chinese Photinia |
| Rutaceae | *Skimmia* japonica | Japanese Skimmia |
| Rosaceae | *Pyracantha* sp. | Firethorn |
| Rosaceae | *Prunus* lusitanica | Portuguese Laurel Cherry |
| Elaeagnaceae | *Hippophae* rhamnoides | Sea Buckthorn |
| Fabaceae | *Medicago* sativa | Alfalfa |
| Elaeagnaceae | *Elaeagnus* × submacrophylla | Ebbing’s Silverberry |

**Table S2.** Complete list of primary and secondary antibodies used, the epitopes they bind, and the studies in which they were described

| Code | Origin | Specificity | Dilution | Supplier | Ref. |
| --- | --- | --- | --- | --- | --- |
| CBM27A | Polyhistidine | 1,4-β-mannans | 10 µg/mL | NZyTech Portugal | [Boraston et al., 2003](https://doi.org/10.1016/S0969-2126(03)00100-X) |
| CBM77A | Polyhistidine | HG | 10 µg/mL |  | [Venditto et al., 2016](https://doi.org/10.1073/pnas.1601558113) |
| CBM44A | Polyhistidine | MLG; cellulose; XG | 10 µg/mL |  | [Najmudin et al., 2006](https://doi.org/10.1074/jbc.M510559200) |
| CBM80A | Polyhistidine | MLG; cellulose; galactomannan; XG | 10 µg/mL |  | [Venditto et al., 2016](https://doi.org/10.1073/pnas.1601558113) |
| CBM21A | Polyhistidine | 1,4-α-glucans | 10 µg/mL |  | [Machovič et al., 2005](https://doi.org/10.1111/j.1742-4658.2005.04942.x) |
| CBM28A | Polyhistidine | Cellulose; XG | 10 µg/mL |  | [Tsukimoto et al., 2010](https://doi.org/10.1016/j.febslet.2010.02.027) |
| CBM66A | Polyhistidine | Fructans | 10 µg/mL |  | [Burne et al., 1987](https://doi.org/10.1128/jb.169.10.4507-4517.1987) |
| BS-400-2 | Mouse | (1,3)-β-D-glucan | 1:50 | BioSupplies | [Meikle et al., 1991](https://doi.org/10.1007/BF00194507) |
| BS-400-3 | Mouse | (1,3;1,4)-β-D-glucan | 1:50 |  | [Meikle et al., 1994](https://doi.org/10.1046/j.1365-313x.1994.5010001.x) |
| BS-400-4 | Mouse | (1,4)-β-mannan; DP = 4 or more | 1:50 |  | [Pettolino et al., 2001](https://doi.org/10.1007/s004250100606) |
| LM1 | Rat | Extensin and certain arabinogalactans | 1:10 | Plant Probes | [Smallwood et al., 1995](https://doi.org/10.1007/bf00203651) |
| LM2 | Rat | β-linked- GlcA in AGP glycan | 1:10 |  | [Møller et al., 2008](https://doi.org/10.1007/s10719-007-9059-7); [Yates et al., 1996](https://scispace.com/pdf/characterization-of-carbohydrate-structural-features-27r5v5bp7l.pdf) |
| LM5 | Rat | (1,4)-β-D-galactan | 1:10 |  | [Jones et al., 1997](https://doi.org/10.1104/pp.113.4.1405) |
| LM6-M | Rat | (1-5)-α-L-arabinan | 1:10 |  | [Cornuault et al., 2018](https://www.biorxiv.org/content/10.1101/161604v1) |
| LM8 | Rat | Xylogalacturonan | 1:10 |  | [Willats et al., 2004](https://doi.org/10.1007/s00425-003-1147-8) |
| LM9 | Rat | Feruloylated-(1-4)-b-D-galactan | 1:10 |  | [Clausen et al., 2004](https://doi.org/10.1007/s00425-004-1309-3) |
| LM10 | Rat | Non-reducing terminal (1,4)-β-D-xylose in xylan backbone | 1:10 |  | [Ruprecht et al., 2017](https://doi.org/10.1104/pp.17.00737); [McCartney et al., 2005](https://doi.org/10.1369/jhc.4b6578.2005) |
| LM11 | Rat | Xylan backbone; (1,4)-β-D-xylan, DP at least 2 | 1:10 |  | [Ruprecht et al., 2017](https://doi.org/10.1104/pp.17.00737); [McCartney et al., 2005](https://doi.org/10.1369/jhc.4b6578.2005) |
| LM13 | Rat | Linearised (1, 5)-α-L-arabinan | 1:10 |  | [Møller et al., 2008](https://doi.org/10.1007/s10719-007-9059-7); [Verhertbruggen et al., 2009](https://doi.org/10.1016/j.carres.2008.11.010) |
| LM14 | Rat | GlcA in AGP glycan | 1:10 |  | [Møller et al., 2008](https://doi.org/10.1007/s10719-007-9059-7) |
| LM15 | Rat | Non-reducing terminal X or GX-motif in xyloglucan | 1:10 |  | [Marcus et al., 2008](https://doi.org/10.1186/1471-2229-8-60); [Ruprecht et al., 2017](https://doi.org/10.1104/pp.17.00737) |
| LM16 | Rat | Processed (1, 5)-α-L-arabinan | 1:10 |  | [Verhertbruggen et al., 2009](https://doi.org/10.1016/j.carres.2008.11.010) |
| LM18 | Rat | HG partially/un-esterified | 1:10 |  | [Verhertbruggen et al., 2009](https://doi.org/10.1016/j.carres.2008.11.010) |
| LM19 | Rat | HG partially esterified | 1:10 |  | [Ruprecht & Pfrengle, 2022](https://doi.org/10.1007/978-1-0716-2148-6_7); [Verhertbruggen et al., 2009](https://doi.org/10.1016/j.carres.2008.11.010) |
| LM20 | Rat | HG; every other GalA esterified | 1:10 |  | [Ruprecht & Pfrengle, 2022](https://doi.org/10.1007/978-1-0716-2148-6_7); [Verhertbruggen et al., 2009](https://doi.org/10.1016/j.carres.2008.11.010) |
| LM21 | Rat | (1,4)-β-D-mannan/galactomannan | 1:10 |  | [Marcus et al., 2010](https://doi.org/10.1111/j.1365-313x.2010.04319.x) |
| LM23 | Rat | Non-reducing terminal (1,4)-β-D-xylose disaccharide in xylan backbone | 1:10 |  | [Pedersen et al., 2012](https://doi.org/10.1074/jbc.m112.396598); [Ruprecht et al., 2017](https://doi.org/10.1104/pp.17.00737) |
| LM24 | Rat | Galactosylated xyloglucan | 1:10 |  | [Pedersen et al., 2012](https://doi.org/10.1074/jbc.m112.396598) |
| LM25 | Rat | Non-reducing terminal X or GX-motif in xyloglucan | 1:10 |  | [Pedersen et al., 2012](https://doi.org/10.1074/jbc.m112.396598); [Ruprecht et al., 2017](https://doi.org/10.1104/pp.17.00737) |
| LM26 | Rat | (1,6)-galactosyl substitution in (1,4)-β-D-galactan | 1:10 |  | [Torode et al., 2018](https://doi.org/10.1104/pp.17.01568) |
| LM28 | Rat | Glucuronoxylan | 1:10 |  | [Cornault et al., 2015](https://doi.org/10.1007/s00425-015-2375-4) |
| JIM13 | Rat | AGP | 1:10 |  | [Yates et al., 1996](https://doi.org/10.1093/glycob/6.2.131) |
| JIM16 | Rat | AGP glycan | 1:10 |  | [Yates et al., 1996](https://doi.org/10.1093/glycob/6.2.131) |
| MAC207 | Mouse | AGP glycan | 1:10 |  | [Pennel et al., 1989](https://doi.org/10.1083/jcb.108.5.1967) |
| INCh1 | Mouse | 1,4-α-glucans | 1:10 | INRAe | [Rydahl et al., 2017](https://doi.org/10.1038/s41598-017-04307-2) |
| RbcL (1) | Rabbit | RbcL | 1:1000 | Abcam |  |
| Lhcb2 | Rabbit | LHCII type II chlorophyll a/b-binding protein | 1:1000 | Agrisera | [Lang et al., 2011](https://pubmed.ncbi.nlm.nih.gov/20960201/) |
| Lhcb4 | Rabbit | CP29 chlorophyll a/b binding protein of plant PSII | 1:1000 |  | [Grimmer et al., 2014](http://journal.frontiersin.org/Journal/10.3389/fpls.2014.00258/abstract) |
| RbcL (2) | Rabbit | RbcL | 1:1000 |  | [Mayfield et al., 2014](http://www.ncbi.nlm.nih.gov/pubmed/24449387) |
| RbcS | Rabbit | RbcS | 1:1000 |  | [Sun et al., 2014](http://link.springer.com/article/10.1007%2Fs11738-014-1648-z) |
| RPE | Rabbit | RPE | 1:1000 |  |  |
| PsbP | Rabbit | Psbp | 1:1000 |  | [Chen et al., 2017](http://www.sciencedirect.com/science/article/pii/S0098847216302519) |
| GLDP | Rabbit | GLDP | 1:1000 |  | [Khoshravesh et al., 2020](https://www.ncbi.nlm.nih.gov/pubmed/31611421) |
| ACT | Rabbit | ACT | 1:1000 |  | [Zheng et al., 2014](http://www.ncbi.nlm.nih.gov/pubmed/24824475) |
| PEPC | Rabbit | PEPC | 1:1000 |  | [Aragón et al., 2013](http://www.ncbi.nlm.nih.gov/pubmed/23959598) |
| RA | Rabbit | RA | 1:1000 |  | [Chen et al., 2013](http://www.ncbi.nlm.nih.gov/pubmed/24349468) |
| PRK | Rabbit | PRK | 1:1000 |  | [Nikkanen et al., 2016](https://www.ncbi.nlm.nih.gov/pubmed/26831830) |
| NTRC | Rabbit | NTRC | 1:1000 |  |  |
| Negative  Rat | Rat | Secondary mAb |  | Jackson ImmunoResearch |  |
| Negative Mouse | Mouse | Secondary mAb |  |  |  |
| Negative Rabbit | Rabbit | Secondary mAb |  |  |  |
| Negative Polyhistidine | Polyhistidine | Secondary mAb |  | Sigma |  |

**Table S3.** Plant material weights required with enzyme concentrations

| Species | Frozen-thawed leaf weight | Frozen-thawed pulp weight | Freeze-dried leaf and pulp weight | Enzyme concentration |
| --- | --- | --- | --- | --- |
| *Solanum* lycopersicum | ~ 219 mg | ~ 81 mg | ~ 30 mg | 1. Celluzyme 0.7T – 0.5 % w/v 2. Pectinex Ultra SP-L – 0.4 % w/v 3. Macerozyme R-10 – 0.2 % w/v 4. Driselase – 0.1 % w/v 5. Xylio Pre – 0.1 % w/v 6. Pectinex Ultra SP-L + Macerozyme R-10 – 0.2 % and 0.1 % w/v 7. Pectinex Ultra SP-L + Celluzyme 0.7T – 0.2 % and 0.25% w/v |
| *Cucumis* sativus | ~ 176 mg | ~ 80 mg |  |  |
| *Lolium* perenne | ~ 149 mg | ~ 73 mg |  |  |

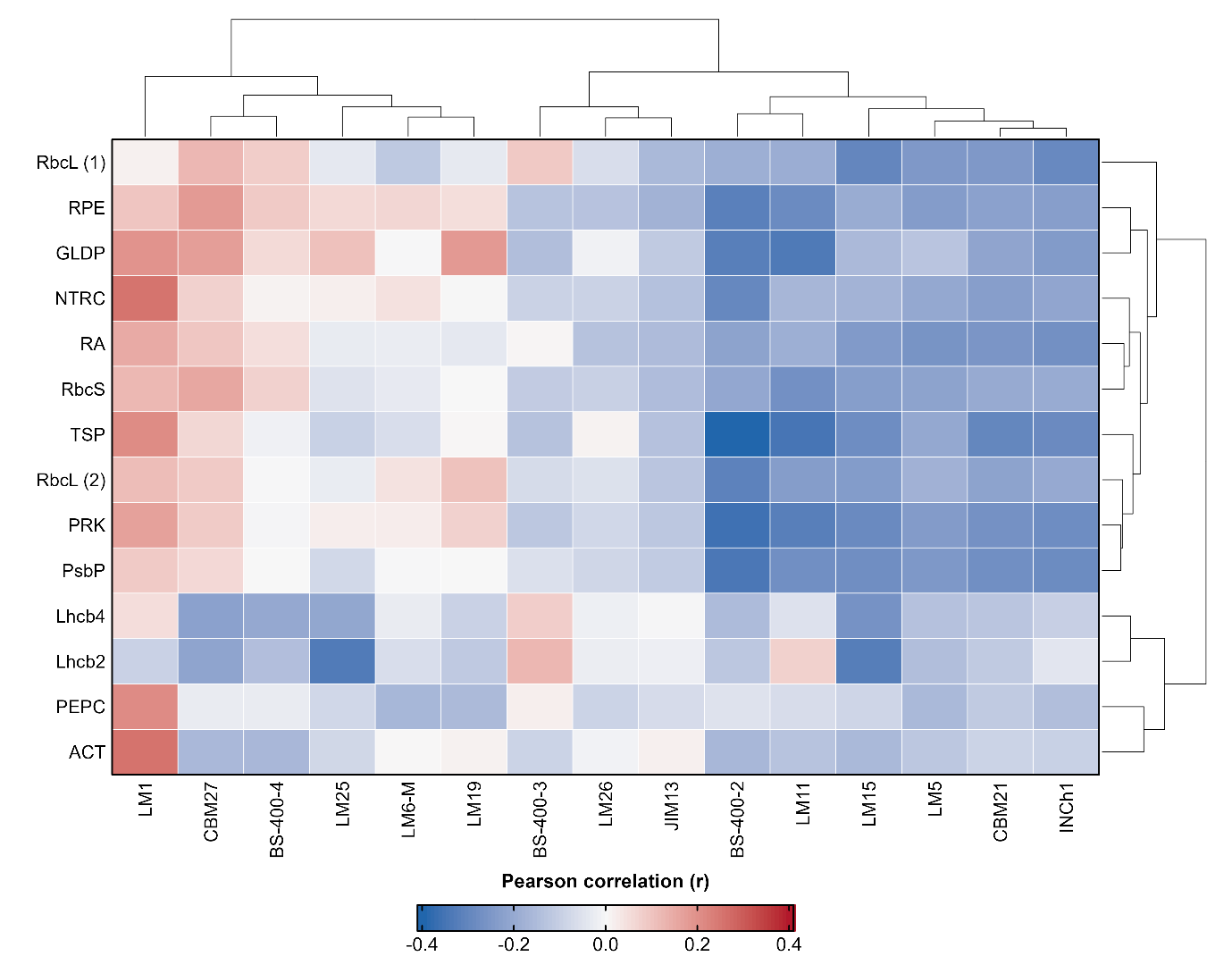

**Fig. S1.** Sensitivity analysis of Pearson correlations between total soluble protein, protein-probe and glycan-probe signals across the plant biomass panel.

The analysis shown in Figure 6 was repeated with total soluble protein (TSP) included as an additional row. Each cell represents the Pearson correlation coefficient (*r*) between mean TSP concentration or mean protein-probe signal intensity and mean glycan-probe signal intensity across 128 plant samples. Red indicates positive correlations, blue indicates negative correlations and near-white cells indicate little or no linear association. Protein probes and TSP were hierarchically clustered using Euclidean distance and complete linkage according to the similarity of their complete glycan-correlation profiles; glycan probes were clustered according to their relationships across the protein probes and TSP. TSP was quantified using the Bradford assay. RbcL (1) and RbcL (2) are Abcam and Agrisera antibodies against the Rubisco large subunit, respectively; RbcS is an Agrisera antibody against the Rubisco small subunit. Further probe identities and specificities are provided in Table 2.

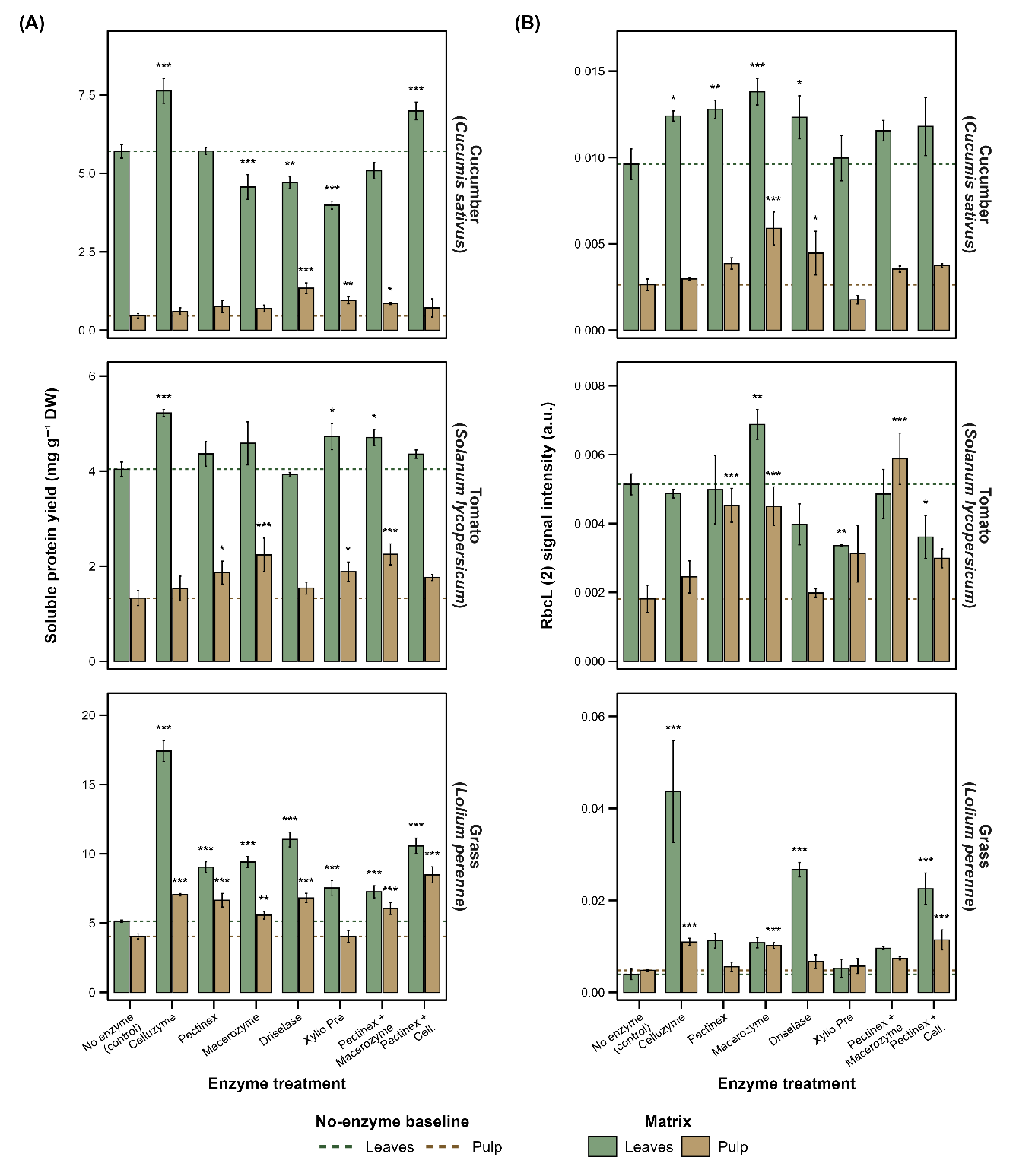

**Fig. S2. Effects of enzyme treatment on total soluble protein yield and RbcL (2) signal in freeze-dried plant biomass.**

(A) Total soluble protein yield and (B) MAPP-derived RbcL (2) signal intensity in leaf and pulp fractions of cucumber (Cucumis sativus), tomato (Solanum lycopersicum) and perennial ryegrass (Lolium perenne). Bars show means ± SD from three biological replicates (n = 3). Green and brown bars represent leaf and pulp fractions, respectively; dotted horizontal lines indicate the corresponding mean no-enzyme controls. Asterisks indicate Dunnett-adjusted differences from the matched species- and matrix-specific no-enzyme control: p < 0.05, **p** < 0.01 and ***p*** < 0.001. RbcL (2) denotes the Agrisera antibody targeting the Rubisco large subunit. Biomass weights and enzyme concentrations are provided in Table 3. DW, dry weight.
